# Progressive protein aggregation in PRPF31 patient retinal pigment epithelium cells: the mechanism and its reversal through activation of autophagy

**DOI:** 10.1101/2021.10.11.463925

**Authors:** Maria Georgiou, Chunbo Yang, Robert Atkinson, Kuan-Ting Pan, Adriana Buskin, Marina Moya Molina, Joseph Collin, Jumana Al-Aama, Franziska Goertler, Sebastian E. J. Ludwig, Tracey Davey, Reinhard Lührmann, Sushma Nagaraja-Grellscheid, Colin Johnson, Robin Ali, Lyle Armstrong, Viktor Korolchuk, Henning Urlaub, Sina Mozaffari-Jovin, Majlinda Lako

## Abstract

Mutations in pre-mRNA processing factor 31 (PRPF31), a core protein of the spliceosomal tri-snRNP complex, cause autosomal-dominant retinitis pigmentosa (adRP). It has remained an enigma why mutations in ubiquitously expressed tri-snRNP proteins result in retina-specific disorders, and so far, the underlying mechanism of splicing factors-related RP is poorly understood. Here, we used iPSC technology to generate retinal organoids and RPE models from three patients with severe and very severe PRPF31-adRP, normal individuals and a CRISPR/Cas9-corrected isogenic control. To fully assess the impacts of *PRPF31* mutations, quantitative proteomics analyses of retinal organoids and RPE cells was carried out showing RNA splicing, autophagy and lysosome, unfolded protein response (UPR) and visual cycle-related pathways to be significantly affected. Strikingly, the patient-derived RPE and retinal cells were characterised by the presence of large amounts of cytoplasmic aggregates containing the mutant PRPF31 and misfolded, ubiquitin-conjugated proteins including key visual cycle proteins, which accumulated progressively with time. Mutant PRPF31 variant was not incorporated into splicing complexes, but reduction of PRPF31 wildtype levels led to tri-snRNP assembly defects in Cajal bodies of PRPF31 patient retinal cells with reduced U4/U6 snRNPs and accumulation of U5, smaller nuclear speckles and reduced formation of active spliceosomes giving rise to global splicing dysregulation. Moreover, the impaired waste disposal mechanisms further exacerbated aggregate formation, and targeting these by activating the autophagy pathway using Rapamycin resulted in reduction of cytoplasmic aggregates and improved cell survival. Our data demonstrate that it is the progressive aggregate accumulation that overburdens the waste disposal machinery rather than direct PRPF31-initiated mis-splicing, and thus relieving the RPE cells from insoluble cytoplasmic aggregates presents a novel therapeutic strategy that can be combined with gene therapy studies to fully restore RPE and retinal cell function in PRPF31-adRP patients.

**Highlights:**

1. *PRPF31* RP mutations lead to formation of insoluble aggregates containing the mutant PRPF31 and misfolded, ubiquitin conjugated proteins including key visual cycle proteins (e.g. RLBP1) in RPE cells, which accumulate progressively with time and affect tight junctions and cell survival.
2. Mutant PRPF31 is predominantly localised in cytoplasmic aggregates of patient specific RPE and retinal cells and is not able to be incorporated into splicing complexes to cause direct mis-splicing.
3. High-throughput quantitative proteomics identifies significantly altered RNA splicing, visual perception, retinoid metabolism, waste disposal and unfolded protein response pathways in patient RPE cells, and autophagy and lysosome, unfolded protein response (UPR) and visual cycle-related pathways in photoreceptor cells.
4. Accumulation of PRPF31 mutant variant as cytoplasmic aggregates reduces wildtype PRPF31 in the nucleus leading to tri-snRNP assembly defects, characterised by accumulation of U5 and reduction of U4/U6 snRNPs in Cajal bodies, altered morphology of nuclear speckles and consequently downregulation of active spliceosomes (B^act^ and C complexes) in PRPF31 patient RPE and retinal cells.
5. Proteomic study of insoluble aggregates identifies other RP-linked splicing factors and multiple key retinal-specific proteins, whose variants are linked to retinitis pigmentosa, within the aggregates of patient RPE cells.
6. PRPF31 patient RPE cells have impaired waste disposal and proteasome mediated degradation, which together with the impaired autophagy pathway, further exacerbate aggregate formation.
7. Phagocytosis of photoreceptor outer segment fragments (POS) shed daily by RPE cells accelerates aggregation of key proteins indicating enhanced cytoplasmic aggregate formation under physiological conditions in patient RPE cells.
8. Activation of autophagy via administration of rapamycin results in reduction of cytoplasmic aggregates in RPE cells, correct localisation of mislocated and misfolded proteins to the nucleus, thereby improving cell survival.

## Introduction

Misfolding, aggregation, and deposition of abnormal proteins is a common hallmark event of multiple neurodegenerative diseases (NDs). Aberrant accumulation of self-aggregating proteins intracellularly and extracellularly causes cellular toxicity due to the formation of insoluble non-native aggregates, which disrupt protein homeostasis and eventually lead to cellular dysfunction or cell death (1). Although protein aggregates differ in protein composition, size, and structure in distinct NDs, they share common cytotoxic effects and accumulate progressively over time (2).

Aggregation and accumulation of misfolded proteins over time is also a common feature of many retinal dystrophies including age-related macular degeneration and retinitis pigmentosa (RP) (3). RP is the most common among all inherited retinal disorders causing blindness, with an incidence of 1 in 4000 people, and more than 1 million affected individuals worldwide (4). The progressive degeneration of photoreceptor cells and retinal pigmented epithelium (RPE) cells are the major pathological events in autosomal dominant RP cases (5), which constitute approximately 43% of all known RP cases.

Up to date, more than 80 genes have been implicated in non-syndromic RP. These include genes that encode retinal-specific proteins such as *Rhodopsin* (*RHO*), the most frequently affected gene in RP, associated with the formation of cellular aggregates (6). Particularly, *RHO*^P23H^, which is the most common mutation in RP, produces an extremely aggregation-prone form of rhodopsin failing to translocate to the plasma membrane to form the visual pigment (3). These aggregates are ubiquitinated and targeted for degradation by the proteasome (6), however, saturation of the proteolytic machinery enhances the accumulation of rhodopsin aggregates (7).

In addition to the genes encoding retinal-specific proteins, mutations in genes encoding pre-mRNA processing factors *(PRPFs)* have been associated with retina-specific diseases, despite their ubiquitous expression (8). PRPFs including PRPF3, PRPF4, PRPF6, PRPF8, PRPF31, and SNRNP200, are all components of the U4/U6.U5 tri-snRNP complex, which is a major constituent of the spliceosome— a large macromolecular complex essential for the catalysis of pre-mRNA splicing (9). Mutations in *PRPFs* constitute 15% of adRP cases and affect the assembly of the spliceosome leading to mis-splicing of genes important for retinal function (10). Published evidence indicates that mammalian cells with *PRPF* mutations are characterised by accumulation of less soluble proteins that are prone to aggregation (11). For example, mutations in *PRPF3* affect the localisation of PRPF3 protein itself leading to the aggregation of misfolded proteins, which triggers the apoptosis of photoreceptor cells (11). However, the role of misfolded aggregates in disease pathogenesis and their association with photoreceptor cell death and/or RPE has not been fully understood.

About 10% of adRP cases are caused by mutations in *PRPF31*, which is an important component of the splicing machinery required for the assembly and stability of tri-snRNPs (12). This type of RP is known as RP11. Using iPSC-based modelling and differentiation to retinal cells, we previously reported the presence of large deposits on the basal side of RP11-RPE cells. Importantly patient specific RPE (but not photoreceptor cells or non-retinal cells) were characterised by the presence of mutant PRPF31 protein, suggesting that RPE cells are the most affected cell type (10). Recently, findings by Diaz-Corrales and colleagues in the Prpf31^p.A216P/+^ mouse model have demonstrated the aggregation of mutant *PRPF31* protein in the cytoplasm of RPE cells (13), accompanied by the overexpression of HSPA4L chaperone (13).

Molecular chaperones act as the initial defence cellular mechanism in response to misfolded proteins damaged by mutations or stress. Under normal conditions, chaperones protect cells by stabilisation of folding intermediates and prevention of protein misfolding and aggregation. However, misfolded proteins that are unable to reassemble correctly are ubiquitinated and targeted for degradation by the proteolytic degradation machinery. In the case where unfolded protein response malfunctions, an intrinsic apoptotic pathway is activated as a secondary response to degrade accumulated proteins. However, dysregulation of autophagy often leads to protein aggregation diseases (14).

In this study, we have used induced pluripotent stem cell-derived RPE (iPSC-RPE) from patients harbouring two different *PRPF31* mutations to investigate their impact on the proteome and phenome of RPE cells. We show that the mutant PRPF31 protein is mis-localised, accumulated, and aggregated in the cytoplasm of RP11-RPE cells in association with misfolded, ubiquitin-conjugated proteins. Importantly, we provide evidence that protein degradation and waste disposal mechanisms are impaired in RP11-RPE cells, leading to the accumulation and deposition of large aggregates, which affect RPE cell survival. Activation of autophagy by Rapamycin enhanced the clearance of cytoplasmic aggregates and improved cell survival.

## Materials and Methods

### Human cell lines

All samples used in this study were obtained with informed consent according to the protocols approved by Yorkshire and the Humber Research Ethics Committee (REC ref. no. 03/362). PRPF31-iPSC lines used in this study were derived from three patient with severe (RP11S1, RP11S2, RP11S3), moderate (RP11M)and very severe (RP11VS) phenotypes as described in our earlier work (10). RP11VS, RP11M and RP11S1 cell lines harbour the same *PRPF31* mutation (c.1115_1125 del11) but vary in the severity of the disease. RP11S2 and RP11S3 patients harbour a different mutation (c.522_527+10del). Crispr/Cas9 corrected (Cas9-RP11VS) and unaffected cell lines (WT1, WT2 and WT3) were used as controls (10).

### iPSC culture

Human iPSCs were cultured on pre-coated with growth factor reduced Matrigel (Corning, 354230) six-well plates. mTeSR™1 (StemCell Technologies, 05850) media supplemented with penicillin/streptomycin (Gibco, 15140) was used on daily basis for the culturing of iPSCs. Passaging of iPSCs was carried out every 4-5 days using Versene (EDTA 0.02%) (Lonza, BE17–771E) solution for 3–5⍰minutes at 37⍰°C. The cells were transferred to fresh pre-coated six well plates in a ratio 1:6. The cells were maintained at 37⍰°C, in a humidified environment, with 5% CO_2_. Freezing of iPSCs was performed using freezing media containing 90% foetal bovine serum (Gibco, 10270) and 10% dimethyl sulfoxide (Sigma, D2650) and 10μM Rock inhibitor (Y-27632, Chemdea, CD0141).

### Differentiation of iPSCs to RPE cells

Control and patient derived iPSCs were grown on Matrigel coated six well plates using mTeSR™1 media. When 80-95% confluency was reached, mTeSR™1 media was replaced with 2ml of differentiation medium (DMEM (High Glucose, 50 μM ß-mercaptoethanol, 1 x MEM NEAA, 20% KOS and 10 mM Nicotinamide) from day 0 to day 7. Subsequently, from day 7 to day 14, nicotinamide was replaced with 100 ng/ml Activin A. Thereafter, nicotinamide was substituted with 3 μM CHIR99021 (Sigma, SML1046) from day 14 to 42. From day 42 onwards until harvesting of the RPE patches, the cells were fed three times a week with differentiation medium containing DMEM (High Glucose), 50 μM ß-mercaptoethanol, 1 x MEM NEAA and 4% KOS. RPE patches were mechanically collected around day 90 using a blade. The collected RPE patches were dissociated in TrypLE (10×) (Invitrogen, USA) for 20⍰minutes at 37°C. The RPE cells were sieved using a 100⍰μm cell strainer and re-plated at 1.5⍰×⍰10^5^ cells per cm^2^ on 12-well plates or on 24 Transwell inserts (GreinerBioOne, 662641).

### Differentiation of iPSCs to Retinal Organoids

iPSCs were dissociated to single cells using Accutase (Gibco, A1110501) and 7,000 cells were seeded into each well of Lipidure-coated 96-well U-bottomed plate (Amsbio AMS LCP-A-U96-6) and cultured in mTeSR-1 medium supplemented with 10 uM ROCK inhibitor Y-27632 (Chemdea, CD0141) to form Embryonic Bodies. After 48 hours, the media was changed to differentiation medium containing 45% Iscove’s modified Dulbecco’s medium (Gibco, 12440–053), 45% Hams F12 (Gibco, 31765–029), 10% KSR (Gibco, 10828–028), 1x GlutaMAX (Gibco, 35050–038), 1% chemically defined lipid concentrate (Thermo, 11905031), 450 μM monothioglycerol (Sigma, M6145), 1x penicillin/streptomycin (Gibco, 15140–122). This was defined as day 0 of differentiation. From day 6 to day 9, 1.5 nM BMP4 (R&D, 314-BP) was added to the differentiation medium. From day 18, the culture medium was changed to 10% FBS (Gibco, 10270–106) in DMEM/F12 (Gibco, 31330–038) supplemented with 1x N2 (Thermo, A1370701), 0.1mM Taurine (Sigma, T8691) and 0.5 uM Retinoic Acid (Sigma, R2625), 0.25 μg/ml Fungizone (Gibco, 15290–02), penicillin/ streptomycin (Gibco, 15140–122) until day 150.

### Western blotting

Cell culture samples were washed three times with PBS and collected. RPE cells were collected using a cell scraper and retinal organoids were manually transferred into 15ml Falcon tubes and precipitated by gravity. Cell culture samples were re-suspended in lysis buffer (25mM Tris-Cl pH 7.5, 120mM NaCl, 1 mM EDTA pH 8.0, 0.5% Triton X100) supplemented with protease inhibitors (Roche 11697498001) and vortexed for 15 minutes at 4°C followed by ultra-sonication three times 5 seconds each (Bradson Sonifier150) to obtain the whole cell lysate. The protein concentration was determined using the Bradford Dye Reagent (Bio-Rad 500-0205). Ten μg of whole cell lysate was applied to SDS-PAGE and transferred to Hybond PVDF membrane (GE Health 15259894). PVDF membranes were blocked for 1 hour in 5% dried skimmed milk in tris-buffered saline and Tween™ 20 solution (TBST). Thereafter, the membranes were incubated overnight at 4°C with primary antibodies in blocking buffer. Washing of the membranes was performed using TBST and the bound primary antibodies were detected using horseradish peroxidase (HRP)-conjugated secondary antibodies. Signal was detected by using SuperSignal West Pico PLUS Chemiluminescent Substrate (ThermoFisher) and visualised by Amersham Imager 600 (GE Healthcare Bio-Sciences AB) imager. The signal intensity was quantified by Image Studio Lite v5.2 (LI-COR Biosciences). Dilution of antibodies is shown in **Table S1.**

### Immunofluorescence (IF) analysis in RPE cells

iPSC-derived RPE cells grown on 24-well PET hanging cell culture inserts (Merck) (pore size 0.4 μm) were fixed with 4% paraformaldehyde (PFA) (Sigma, 47608) for 20⍰minutes at room temperature. Before blocking, the pigmentation of RPE cells was removed using Melanin Bleach Kit (Polysciences), followed by 3 washes with PBS. RPE cells were blocked in PBS supplemented with 10% donkey serum and 0.3% Triton-X100 (Sigma, T8787) to permeabilised the cells, for 1 hour at room temperature. For p62 and LC3 methanol fixation was used instead and no bleaching prior to immunostaining was carried out. The RPE cells were incubated with primary antibodies overnight at 4°C **(Table S1).** Following three washing with PBS, RPE cells were incubated with secondary antibodies **(Table S1)** diluted in antibody dilution (PBS, 1% donkey serum, and 0.1% Triton X-100) and stored overnight at 4°C. Nuclei were stained with Hoechst (Life Technologies, UK). Then, RPE cells were mounted with Vectashield and sealed with a coverslip. RPE cells were imaged using the Axio Imager upright microscope with Apotome structured illumination fluorescence using 20x objective, 40x and 63x oil objectives (Zeiss, Germany). Images were presented as a maximum intensity projection and adjusted for brightness and contrast in Adobe Photoshop (Adobe Systems).

### Fluorescence in situ hybridisation

Fluorescence *in situ* hybridisation (FISH) was performed using probes labelled with Alexa 647 at the 5’-end. The sequences of probes and the RNA-FISH method were as previously described (15, 16). After FISH, cells were immunostained for coilin and prepared for confocal microscopy as described above. Quantification of mean intensities was performed with the ImageJ/Fiji software from four independent measurements.

### Transmission electron microscopy (TEM)

Trans-wells of RPE cells were washed with PBS and then fixed with 2% gluteraldehyde in 0.10M sodium cacodylate buffer. The samples were further fixed in 1% osmium tetroxide, dehydrated in gradient acetone, and embedded in epoxy resin. Sections of 70 nm thickness were picked up on coper grids, stained with uranyl acetate and lead citrate and imaged using a Philips CM100 transmission electron microscope with high-resolution digital image capture.

### Proteasome Activity Assay

RPE cells from control and patient cell lines were washed with PBS and collected using a cell scraper. Pellets were resuspended in lysis buffer containing 0.5% NP-40 supplemented in distilled water and incubated on ice for 30 minutes followed by centrifugation at 13,000 x g for 20 minutes at 4°C. The supernatant was collected and analysed by Pierce BCA Protein Assay kit (Pierce, ThermoFisher Scientific) to measure the protein concentration. Proteasome substrate Bz-VGR-AMC (BW9375, Biomol International) was used to measure the trypsin like activity of the proteasome. As a control, proteasome inhibitor, MG132, was used. The chemotrypsin like activity was measured at excitation/emission wavelength of 360 nm/460 nm, respectively, using a Varioskan LUX multimode Microplate reader (ThermoFisher Scientific).

### Isolation of insoluble fractions in RPE cells

RPE cells were washed with PBS and collected using a cell scraper. Cell pellets were lysed using lysis buffer containing 10 mM Tris-HCL (Sigma, 1185-53-1), pH 7.5, 5 mM EDTA, 1% NP-40 (Sigma, 127087-87-0), 0.5% deoxycholate, 150 mM NaCl (Sigma, 7647-14-5) and 1 complete ULTRA tablet (EDTA – free protease inhibitor) (Sigma, 06 538 282 001). The lysates were incubated on ice for 15 minutes followed by vortexing at 4°C for 15 minutes. Thereafter, the lysates were sonicated and centrifuged for 15 minutes at 13,000 x g. The generated supernatant was transferred in a fresh tube and labelled as the soluble fraction. The remaining pellets were mixed with 20μl of a lysis buffer (60 mM Tris-HCL (Sigma, 10812846001) pH 7,2% SDS and, 2.5% 2-Mercaptoethanol (Sigma, 60-24-2), and sonicated. Thereafter, the samples were centrifuged at 16000g for 20 minutes at 4°C and the collected supernatant was labelled as the insoluble fraction.

### Quantification and statistical analysis

Statistical analysis was perfumed using Prism (GraphPad, USA). Comparisons between variables and statistical significance between groups were performed using ANOVA and Two-tailed Student’s t-test. Error bars represent standard error of the mean (SEM) unless indicated otherwise. Statistical significance was established as indicated by asterisks *p< 0.05, **p<0.01, *** p<0.001, and **** p<0.0001.

### Treatment of RPE cells with Photoreceptor Outer Segments (POSs)

RPE cells were treated with POS or POS-FITC (20 POS /cell), after cooling of RPE cells for 30 minutes at 17 °C. After incubation of RPE cells with POS or POS-FITC, media was removed and replaced with POS suspension immediately, following by incubation at 17°C for another 30 minutes. This was performed to maximise binding of POSs in RPE cells. Following incubation for 30 minutes, media was aspirated and replaced with fresh warmed media to remove unbound POS. RPE cells kept in a humidified incubator at 37°C with 5% CO_2_. RPE cells were collected at 0, 48, 96, 144 hours post-POS feeding. The RPE cells were washed with PBS twice and then fixed with 4% PFA before immunohistostaining.

### TMT labelling for mass spectrometry

Total cell lysates were prepared from RP11 and control retinal organoid or RPE cells using Pierce Mass Spec Sample Prep Kit (Thermo Scientific). Lysates were diluted to 120 μl and sonicated. Protein concentrations were determined using the Pierce BCA protein assay kit (Thermo Scientific) and 100 μg of the total proteins from each cell line were labelled with 6-plex isobaric tandem mass tag (TMT6) reagents (Thermo Scientific) following the manufacturer’s instruction. To this end, samples were reduced by the addition of tris(2-carboxyethyl)phosphine, alkylated with iodoacetamide and acetone precipitated. Protein pellets were resuspended in 50mM triethyl ammonium bicarbonate (TEAB) buffer and were digested with trypsin overnight at 37°C. For retinal organoid samples, the patient RP11S1, RP11M, RP11S2 lines and control WT2, WT3 samples were, respectively, labelled with TMT6-128, TMT6-129, TMT6-130 and TMT6-126, TMT6-127 reagents for 1 hour at room temperature. For RPE samples, the patient RP11S1, RP11VS, RP11S3 lines and control WT1 samples were, respectively, labelled with TMT6-127, TMT6-128, TMT6-129 and TMT6-126 reagents. Proteomics data from RP11VS and Cas9-RP11VS RPE cells and retinal organoids obtained in our early study were also included for analysis (10). Reactions were quenched by 5% hydroxylamine for 15 minutes. Fifty micrograms of TMT-labelled peptides from RP11 and control cells were combined and cleaned up using C18 spin columns (Harvard Apparatus), dried by SpeedVac (Eppendorf) and subjected to peptide pre-fractionation using high-pH reversed-phase chromatography. After constituting the dried, TMT-labelled peptides in 100 μl buffer A (10mM NH4OH), fifty microliters of peptide mixtures were injected into an XBridge BEH C18 HPLC column (150mm × 1 mm ID, 3.5 μm; Waters) and separated in 80 fractions using a gradient of buffer B (10mM NH4OH, 80% acetonitrile) over 90 min. Collected fractions were combined into 20 fractions, dried and resuspended in 20 μl of 0.1% trifluoroacetic acid (TFA) for mass spectrometry analysis.

### LC-MS/MS analysis

Peptides in each fraction were analysed in three replicates using either an Orbitrap Fusion or a Q Exactive HF-X mass spectrometer (Thermo Fisher Scientific), both coupled with an UltiMate 3000 RSLCnano HPLC system (Thermo Fisher Scientific), as previously described (10).

### Data processing

MS/MS spectra were searched against a Swiss-Prot human database containing 20,341 reviewed protein entries using Mascot algorithm (Matrix Science) via Proteome Discoverer 2.2 (PD, Thermo Fisher Scientific) and were processed as previously described (10). At least two quantifiable unique peptides in each replicate were required for protein quantification. Protein ratios were log transformed and then median normalised in Perseus. These data were combined with our previous proteomic data from RP11VS and its Cas9-corrected isogenic control (10). The reported RPll/control ratios are the average of at least two replicates. To identify the differentially expressed proteins, those proteins with mean a log2 fold change (LFC) less than −0.5 or greater than +0.5 were defined as regulated. Gene Ontology (GO) enrichment analyses were carried out by Metascape (p-value cutoff 0.02) (17) or by the Perseus software version 1.6.2.2 with a Benjamin–Hochberg FDR 2% (18).

## Results

### Cytoplasmic mislocalisation of the mutant PRPF31 protein and altered morphology of nuclear speckles in RP11-retinal cells

Given the ubiquitous expression of PRPF31, the specificity of RP mutations to the retinal tissue has remained an enigma for the pathogenesis mechanism of FRPFs-related RP disease. In earlier work we have compared PRPF31 expression in various cell lines derived from RP patients in Western blots and shown decreased PRPF31 expression and the robust expression of long mutant form of PRPF31 only in RP11-RPE cells (10). Herein, to assess the localisation of the wild-type and mutant forms of PRPF31 in detail, we first performed immunofluorescence analysis in three RP11-RPE cells using an antibody against the N-terminus of PRPF31 that detects both the wild-type and mutant proteins. Clear differences were observed with PRPF31 being localised in the nucleus of control RPE cells and predominantly in the cytoplasm of RP11-RPE cells **(Figure 1A, S1A).** Additionally, the tight junctions stained with ZO1 antibody were disrupted in RP11-RPE cells compared to the control cells **(Figure 1A, S1A),** corroborating previously observed tight and adherens junction associated protein modifications in the retina of animal model of autosomal dominant RP (adRP) (19).

**Figure 1.**
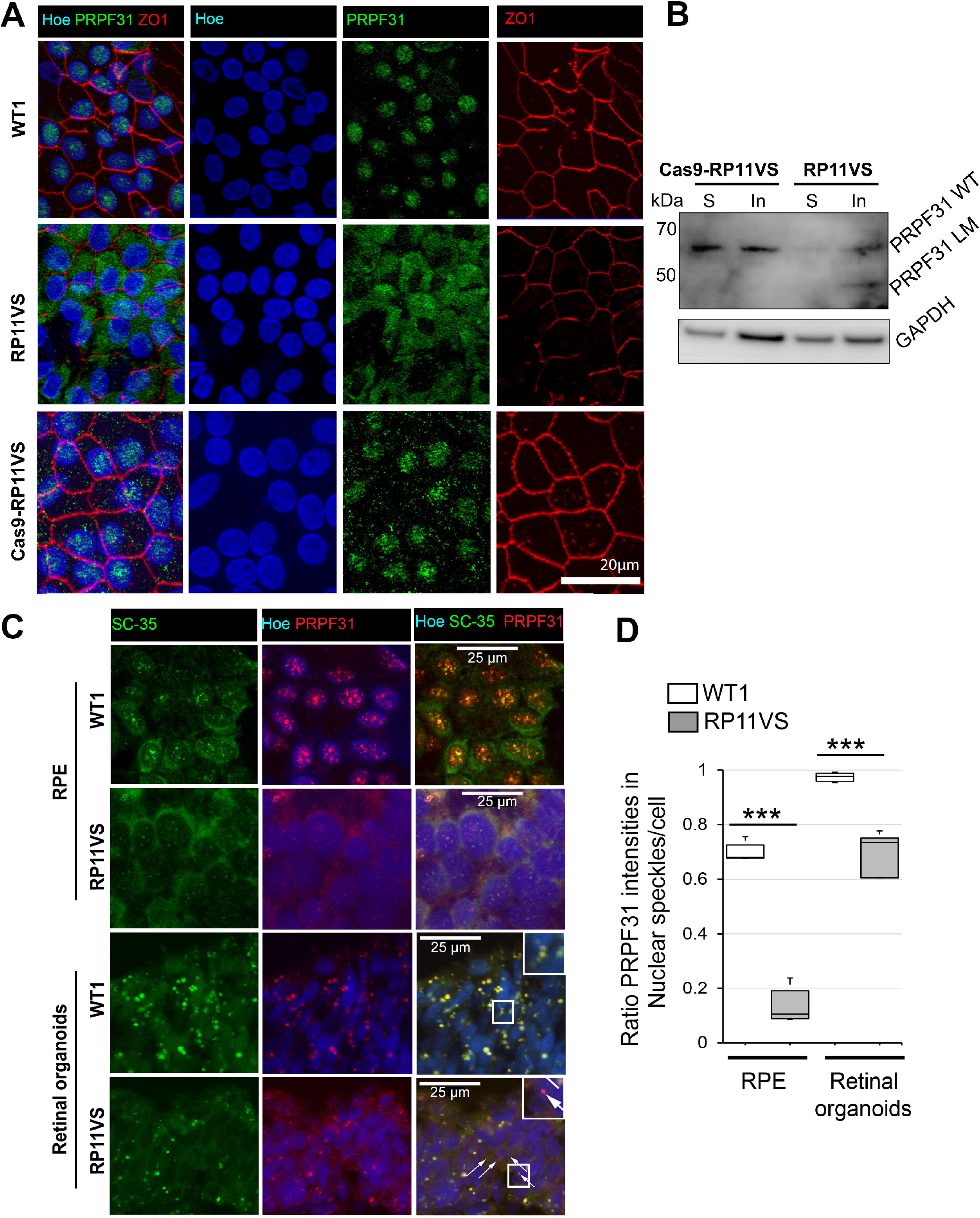
Cytoplasmic localisation of PRPF31 in patient RP11-RPE cells and altered morphology of nuclear speckles and reduced localisation of nuclear PRPF31 in splicing speckles of RP11-RPE and retinal organoids. **A)** Control and RP11-RPE cells were immunostained with an anti-PRPF31 N terminus (green) and ZO1 (red) antibody. Cell nuclei were stained with Hoechst. Immunofluorescence analysis showed localisation of PRPF31 protein mainly in the nucleus of WT1 and RP11VS-Cas9 RPE cells in speckle-like structures, whereas in RP11VS RPE cells, PRPF31 protein is predominantly located in the cytoplasm in an aggregate like pattern. Scale bars: 20 μm. B) Western blot showing the expression of WT and mutant PRPF31 protein in the soluble and insoluble fractions of RP11VS-Cas9 and RP11VS RPE cells with PRPF31 N terminal antibody. GAPDH was used as a loading control. **C)** Co-staining of RPE and retinal organoids with SC35 (SRSF2), a marker for nuclear speckles, and PRPF31 showing altered morphology of nuclear speckles and localisation of PRPF31 in RP11VS cells. Cell nuclei were counterstained with DAPI. Arrows indicate accumulation of mislocalised PRPF31 in the cytoplasm. Scale bars: 25 μm. D) Quantification of signal intensities showing that PRPF31 mislocalisation is more prominent in RPE cells. Representative images from at least three independent experiments are shown.

To distinguish localisation of PRPF31 variants in soluble and insoluble fractions, a Western blot of the two fractions was performed, revealing a dramatic decrease in the amount of PRPF31 in the soluble fraction of RP11-RPE compared to control cells **(Figure 1B).** Notably, whilst the wild-type PRPF31 protein could be detected in the insoluble fraction of both control and RP11 RPE cells, the long mutant isoform was only detected in the insoluble fraction of RP11-RPE suggesting that a significant fraction of cytoplasmic aggregates in RP11-RPE is composed of the mutant PRPF31 **(Figure 1B).** It is known that a large fraction of pre-mRNA splicing occurs co-transcriptionally in decompacted chromatin regions at the periphery or within interchromatin granule clusters, also known as nuclear speckles (20). Nuclear speckles are enriched with splicing factors and thought to be the site for storage of spliceosomal complexes and for post-transcriptional splicing (21) (20). To examine the localisation of PRPF31 to speckles and the effect of *PRPF31* mutations on the morphology of speckles, we co-stained control and RP11-RPE cells and retinal organoids with SC35 (a speckles specific marker) and PRPF31 antibodies. Confocal microscopy revealed large nuclear speckles in control-RPE cells where PRPF31 was localised **(Figure 1C).** In contrast, RP11-RPE cells showed significantly smaller nuclear speckles with reduced signal intensity where only a small fraction of PRPF31 was localised **(Figure 1C, D).** Similar co-staining in retinal organoids revealed large speckles in control photoreceptor cells that significantly decreased in size in RP11-retinal organoids. Moreover, whilst wild type PRPF31 was completely localised to nuclear speckles, we noticed mislocated PRPF31 in the cytoplasm of RP11 retinal organoids **(Figure 1C).** Quantification of signal intensities showed that RPE was the most affected retinal cells for PRPF31 mislocalisation **(Figure 1D).**

Together our data indicate predominant cytoplasmic localisation of PRPF31 isoforms in the RP11-RPE cells, with the long mutant isoform localised in the insoluble protein fraction. Furthermore, the localisation of PRPF31 to nuclear speckles, which are essential for splicing activity, is significantly reduced in both RP11 RPE and retinal organoids.

### Impaired tri-snRNP assembly in Cajal bodies and reduced levels of active spliceosomes in RP11-RPE cells and retinal organoids

Since PRPF31 is an integral component of the U4/U6.U5 tri-snRNP, we next sought to analyse the effect of RP11 PRPF31 variants on the tri-snRNP formation in patient-derived retinal cells. The assembly and maturation of tri-snRNPs occur in Cajal bodies, and defects in the assembly or release of tri-snRNPs results in the accumulation of incomplete snRNPs in Cajal bodies and consequently stalling of spliceosomes at complex A (15, 22). Thus, we performed RNA-FISH using fluorescent probes against U4, U6 and U5 snRNAs followed by immunostaining for coilin, a Cajal body marker. Confocal microscopy and quantification of the fluorescent signals overlapping with Cajal bodies or the nucleoplasm revealed significant reduction of U4 and U6 levels and dramatic accumulation of U5 in Cajal bodies of RP11 photoreceptor cells compared with the controls demonstrating a defect in tri-snRNP formation in these cells **(Figure 2A-D).** These results are also consistent with the reduced size and intensity of nuclear speckles in RP11 retinal organoids and RPE cells. Due to very small Cajal bodies in RPE cells, a reliable co-localisation study of snRNAs was not possible in these cells (data not shown).

**Figure 2.**
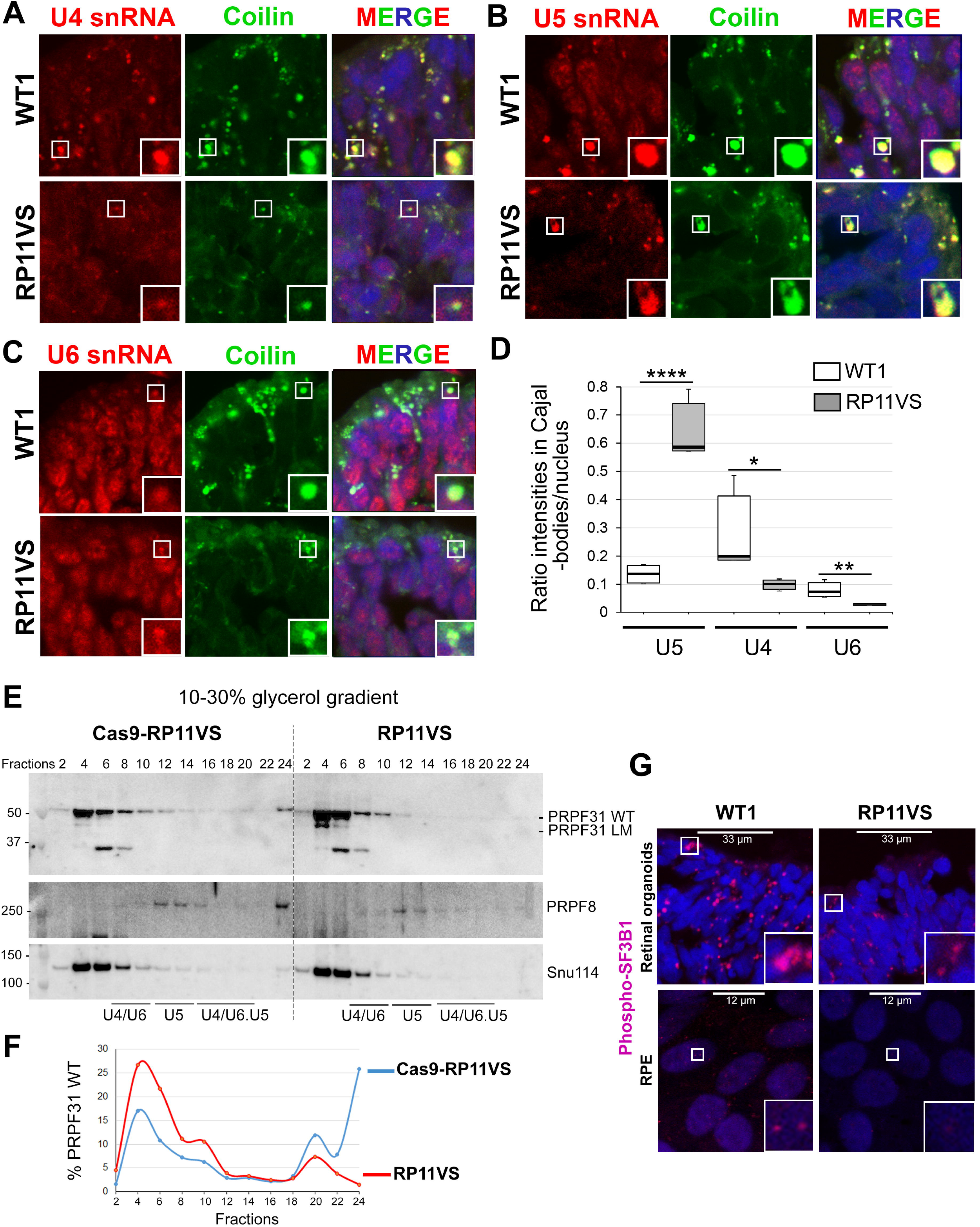
PRPF31 RP mutations lead to defects in the assembly of tri-snRNPs in Cajal bodies, and reduce formation of active spliceosomes (B^act^, C complexes) in RPE and photoreceptor cells. **A-C)** Confocal microscopy analyses of RNA-FISH labelling for U4 (A), U5 (B), and U6 (C) snRNAs (red) in Cajal bodies (anti-coilin, green) in WT1 and RP11VS retinal organoids. D) Quantification of the mean intensity ratios of accumulation of snRNAs in Cajal bodies in RP11VS and WT1 RPE cells. PRPF31 RP mutations lead to the significant accumulation of U5 snRNA and reduction of U4/U6 snRNAs in Cajal bodies. ***P values < 0.001, **P values < 0.01, *P values < 0.05. E) Western blots of even-numbered glycerol gradient fractions of Cas9-RP11VS and RP11VS-RPE cells showing accumulation of PRPF31 wildtype (WT) and long mutant isoform (LM) in the top fractions in RP11VS RPE cells. F) Quantification of the percentage of WT PRPF31 in gradient fractions in Cas9-RP11VS and RP11VS-RPE cells. G) Staining of retinal organoids and RPE cells with anti-phosphorylated SF3B1 antibodies showing reduced levels of active spliceosomes (B^act^ and C complexes) in both RP11VS RPE and photoreceptor cells.

We next analysed formation of spliceosomal subcomplexes by fractionation of RPE and retinal organoids cell extracts on glycerol gradients **(Figure 2E, S2).** Northern blotting of U snRNAs revealed increased levels of snRNAs in the top fractions (3 and 5) for RP11-RPE cells compared with the Cas9-corrected isogenic control **(Figure S2).** Notably, by comparing the snRNA profile of control RPE with that of control retinal organoids, lower amounts of tri-snRNPs in RPE cells were evident. For RP11 retinal organoids, we failed to obtain a reasonable quality of snRNA signals on Northern blots, presumably due to high RNA degradation in the sample that could not be suppressed. Western blotting of even gradient fractions of RPE cells for tri-snRNP proteins PRPF31, PRPF8 and Snu114, showed a significant increase for slow sedimenting PRPF31 in the top fractions (4 and 6) and altered sedimentation pattern of PRPF31-containing spliceosomal complexes in the bottom fractions **(Figure 2E-F).** Importantly, a band corresponding to the PRPF31 long mutant form was detected in fraction 4 of the RP11-RPE indicating that the mutant protein is not integrated into the spliceosomal snRNPs, and thus the mutant PRPF31 cannot directly perturb splicing by inhibition of spliceosome assembly **(Figure 2E).**

To examine the effect of defects in tri-snRNP assembly and changes in the morphology of nuclear speckles on the formation of active spliceosomes in RP11 cells, we next stained RPE and retinal organoids with an anti-phosphorylated SF3B1 antibody. The U2-specific protein SF3B1 is phosphorylated in activated (B^act^) or catalytically active (complex C) spliceosomes, thus can be used as a marker for detection of active spliceosomes (20). Control retinal organoids displayed large foci enriched with p-SF3B1 indicative of the sites of active splicing in photoreceptor cells **(Figure 2G).** These foci were prominently reduced in both size and signal intensity in RP11 organoids demonstrating reduced splicing activity in patient RP11 photoreceptors. We also stained RPE cells with p-SF3B1 antibody and detected a decrease for p-SF3B1 staining in RP11 RPE compared with the control cells **(Figure 2G).**

Altogether, these results show impaired assembly of tri-snRNPs in Cajal bodies of RP11 retinal cells leading to the reduced amounts of active spliceosomes. Furthermore, the mutant PRPF31 protein does not incorporate into snRNP complexes in the nucleus and consequently cannot directly disrupt splicing, but its prominent aggregation in the cytoplasm concomitant with the reduction of wildtype PRPF31 in the nucleus leads to tri-snRNP assembly defects. This in turn can have global impact on the splicing of genes essential for the structure and function of retinal and RPE cells.

### Quantitative proteomics reveals major pathways affected in patient RP11 retinal organoids and RPE cells

#### a. RPE cells

To identify the difference in translational profiles between RP11 patient cells and unaffected controls, high-throughput quantitative proteomics by TMT-labelling of peptides extracted from each cell line and mass spectrometry analysis was performed. RPE cells derived from control (WT1, Cas9-RP11VS) and RP11 patients (RP11S1, RP11S3, RP11VS) identified a total of 5310 proteins of which 1304 proteins were differentially expressed **(Table S2).** Gene ontology (GO) enrichment analyses analysis revealed amongst others enrichment of the DE proteins in RNA splicing, the spliceosome complex, retinoid metabolic process and visual perception, and protein folding pathways **(Table S2).** Of the 90 differentially expressed proteins in the RNA splicing pathway, Fused in Sarcoma (FUS) was the most downregulated protein **(Figure 3A).** Western blot and immunofluorescence analysis corroborated these results and moreover showed that FUS was predominantly localised in the cytoplasm of RP11-RPE in an aggregate-like pattern, unlike control cells where expression was nuclear **(Figure 3B-C).** The mis-localisation of FUS from the nucleus to the cytoplasm has been reported previously by other studies and it has been associated with amyotrophic lateral sclerosis (ALS) (23).

**Figure 3.**
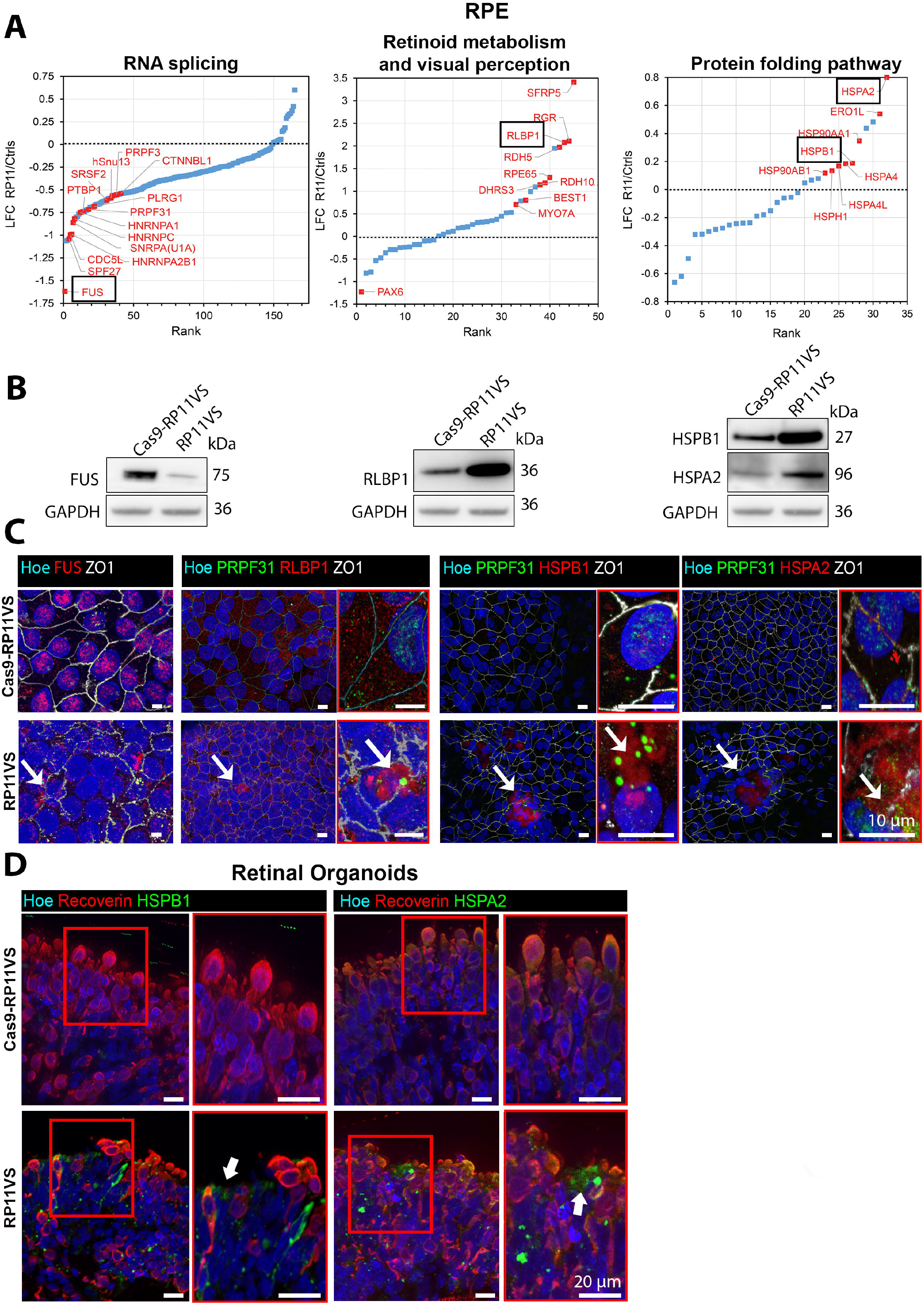
GO analysis reveals enrichment of proteins involved in RNA splicing, retinoid metabolism and visual perception, and protein folding pathway in RP11-RPE cells. **A)** GO analysis shows DE proteins involved in RNA splicing, retinoid metabolism and visual perception, and in protein folding pathway, highlighting with black circle FUS protein, which is the most downregulated protein in RP11-RPE cells from the RNA splicing pathway, RLBP1, and two HSPs; HSPB1, HSPA2. B) Western blot showing downregulation of FUS in RP11-RPE cells and upregulation of RLBP1, HSPB1, HSPA2 in RP11-RPE cells. GAPDH was used as a loading control. C) Immunostaining of RP11-RPE and control RPE cells with FUS (red), RLBP1 (red), HSPB1 (red) and HSPA2 (red). ZO1 (white) was used to define the tight junctions of RPE cells. Fus is located in the cytoplasm of RP11-RPE cells (white arrow), whereas, in control RPE cells Fus is expressed in the nucleus. RLBP1 and HSPs (HSPB1 and HSPA2) are accumulated in the cytoplasm of RP11-RPE cells in the form of aggregates (shown by white arrow) but not in control RPE cells. Magnified images show the PRPF31 immunostaining (shown by white arrow) in RLBP1 and HSPB1 and HAPA2 aggregates (shown by white arrow). Cell nuclei were counterstained with Hoechst. Scale bars: 10 μm. Data obtained from RPE cells at week 12 of differentiation. D) Aggregation of HSPs in RP11 retinal organoids. Co-staining of Cas9-RP11VS and RP11VS retinal organoids with HSPB1 (green) or HSPA2 (green) with Recoverin (red) showing aggregation of HSPs between Recoverin positive cells. Magnified images show co-localisation of HSPs with Recoverin in an aggregate-like form in RP11VS retinal organoids only (shown by white arrows). Cell nuclei were counterstained with Hoechst. Scale bars: 20 μm. Data obtained from retinal organoids day 210 of differentiation.

Nineteen proteins involved in visual perception and 18 proteins in the retinoid metabolic pathway were differentially expressed, and of these the most upregulated protein was RLBP1 **(Figure 3A, Table S2).** RLBP1 is a soluble retinoid carrier, which plays an important role in the regeneration of 11-*cis*-retinol during the visual cycle (24). The upregulation of RLBP1 protein in RP11-RPE cells was further confirmed by Western blotting and by immunofluorescence analysis, showing accumulation of RLBP1 in aggregate-like pattern **(Figure 3B-C).** In areas containing these large aggregates we also detected disruptions of the tight junctions as well as lack of nuclei **(Figure 3C).** Notably, PRPF31 was also associated with the RLBP1 immunostained aggregates in RP11-RPE cells. Upregulation of visual cycle genes in RPE cells has been associated with associated accumulation of retinoid by-products in the aged RPE cells (25). These results suggest that RP11-RPE cells might develop similar features to aged RPE.

Twenty-three differentially expressed proteins were enriched in the protein folding pathway **(Table S2).** Moreover, two molecular chaperones involved in the UPR pathway **(Figure 3A),** namely HSPB1 and HSPA2 which are activated in response to misfolded proteins to restore protein conformation and to prevent protein aggregation, were upregulated in RP11-RPE cells **(Figure 3A).** Upregulation of HSPs in RP11-RPE cells was confirmed by Western blotting **(Figure 3B),** and thereafter by immunofluorescence analysis **(Figure 3C),** demonstrating HSPB1 and HSPA2 expression in an aggregate-like pattern predominantly located in areas lacking nuclei and characterised by disrupted ZO1 staining **(Figure 3C).** Strikingly and in accordance with results obtained in RPE cells, we also noticed the presence of aggregate-like structures of HSPB1 and HSPA2 between patient RP11 photoreceptors **(Figure 3D),** suggesting a similar mechanism of protein misfolding and UPR activation in RP11-photoreceptors.

#### b. Retinal organoids

Quantitative proteomics of retinal organoids differentiated from an isogenic control and unaffected iPSCs (Cas9-RP11VS, WT2 and WT3) were compared to RP11-retinal organoids (RP11VS, RP11S1, RP11M and RP11S2) (10), resulting in identification of 7446 proteins, of which 1596 proteins were differentially expressed (DE) after applying a LFC cut-off of +/− 0.5 **(Table S3).** Gene ontology enrichment analyses revealed an enrichment for proteins involved in several pathways; however, in view of PRPF31 aggregation in the cytoplasm and main function of photoreceptors in light transduction we focused on proteins involved in endoplasmic reticulum (ER) lumen, autophagy and lysosome, retinoid metabolism and visual perception, response to ER stress and unfolded protein response (UPR) **(Table S3).**

Since PRPF31 is a core splicing factor, we first analysed changes for proteins of the RNA splicing pathway. A total of 258 splicing proteins were identified **(Figure S3A, Table S3),** among which PRPF31 itself **(Figure S3A, C)** and several nuclear ribonucleoproteins (hnRNPD, hnRNPK, hnRNPL and hnRNPR) showed mild to strong downregulation. HnRNPs play role in multiple aspects of nucleic acid metabolism regarding transcription, alternative splicing, mRNA stability, transportation, and translation (26, 27), and their reduced expression indicates that various aspects of mRNA metabolism in RP11 organoids are disrupted from transcription to maturation, translation, and turnover. The most downregulated splicing factors in RP11 retinal organoids were NOVA1 and PTB2, which function as splicing repressors and activators (28). NOVA1 and PTBP2 are highly expressed in the brain and involved in neuron specific alternative splicing (29). Since retina is part of the central nervous system, the significant downregulation of NOVA1 and PTBP2 may imply an important role for these splicing factors that deserves further investigation in the pathogenesis of RP11.

GO cellular compartment (GOCC) enrichment analysis also identified 12 DE proteins in the autophagy pathway **(Figure S3B, Table S3).** Autophagy is a lysosomal degradation process for the clearance of damaged organelles and unfolded proteins to maintain cellular homeostasis. During autophagy, cytoplasmic cargos are enclosed in a double-membraned autophagosome followed by fusion with the lysosome and final degradation. Autophagy procedure basically includes three phases: autophagosome formation, autophagosome–lysosome fusion, and degradation (30). First, Rab GTPase family members Rab1A, Rab12 and Rab33B, as well as Rab23 interact closely with one another to form a functional cluster. Dynamic membrane trafficking events are essential for autophagy, thus vesicular trafficking proteins, Rab GTPases play important roles in various steps of autophagy process including vesicular formation, transport, tethering and fusion (31). Interestingly, Rab27a and Rab38 levels show more than a 3-fold increase in RP11 organoids **(Figure S3B, Table S3).** The elevated levels of these proteins in unprenylated form have been implicated in choroidermia, a retinal degeneration disease leading to blindness in late adulthood (32). Moreover, increased levels of Rab27a and Rab27b proteins, which are localized to autophagosomes under stress conditions have been associated with multiple neurodegenerative diseases such as Parkinson’s disease and shown to be essential for the clearance of α-Synuclein by promoting autophagy (33). The upregulation of these three Rab proteins (1.2 times for Rab1A, Rab12, 33B and 1.5 times for Rab23) in the RP11-retinal organoids suggests an enhanced vesicular trafficking and the stimulation of autophagy.

Along with the enrichment of proteins related to autophagy, GOCC enrichment analysis identified 142 DE proteins associated with the lysosome as one of the most enriched compartments, with most of the proteins in this pathway being significantly upregulated **(Table S3).** Amongst these proteins, increased expression of two lysosomal-associated membrane proteins LAMP-1 and LAMP-2 in the RP11-retinal organoids was also confirmed by Western blotting **(Figure S3C).** LAMP-1 and LAMP-2 are key components of the lysosomal membrane that account for around 50% of all the proteins (34). These two proteins facilitate the lysosomes-autophagosome fusion process (35, 36), and their upregulation in RP11-retinal organoids could signal an expansion of the lysosomal complex that is needed to cope with autophagic clearance of mis-spliced and/or misfolded proteins ensuing from the global spliceosome dysregulation (10). In accordance with autophagy stimulation, GOCC enrichment analysis identified 100 DE proteins localised to the ER lumen as one of the most significantly enriched compartments **(Figure S3D, Table S3),** including DNAJB9, DNAJC3, P4HB, HSPA5, as well as EDEM3 and ERLEC1 involved in ER-associated degradation. Notably, most of these DE proteins showed a trend for upregulation, and THBS1 involved in ER stress response was the most upregulated protein, suggesting an elevated unfolded protein response (UPR) activity in RP11-retinal organoids. The endoplasmic reticulum (ER) is an important organelle responsible for synthesis, folding, modification, and transport of proteins. ER stress occurs due to imbalance between inward flux of nascent polypeptides and the protein-folding machinery. This leads to activation of the UPR pathway to adjust ER folding capacity and maintain protein homeostasis (37). Consistently with this, GO biological processes (GOBP) enrichment analysis by Metascape further identified 26 DE proteins in the UPR pathway **(Table S3, Figure S3E).** Together these data suggest that PRPF31 mutation-induced splicing defects may promote protein misfolding (including PRPF31 itself), resulting in activation of UPR and recruitment of heat shock proteins to the aggregates.

GOBP enrichment analysis identified 31 DE proteins involved in visual perception as one of the most enriched compartments **(Figure S3F, Table S3)** including RLBP1, RDH11 and RDH5, which are important proteins involved in the generation of the visual pigment and hence the maintenance of vision. Dysregulated expression of proteins involved in the visual cycle suggest changes in retinoid by-products in retinal organoids, which have been associated with retinal degeneration (25) and deserve further investigations.

### Proteomic analysis of cytoplasmic aggregates found in RP11-RPE cells

Our studies have shown the presence of wild type and mutant PRPF31 in the insoluble aggregates of RP11-RPE cells **(Figure 1B).** To assess in detail the composition of the insoluble fraction we performed comparative proteomic analysis of the insoluble fractions prepared from control and RP11-RPE cells separated by SDS-PAGE and analysed by mass spectrometry. Proteomic analysis detected a total of 4061 proteins, of which 934 were DE in patient RP11-RPE cells (LFC cut off value of 1) **(Figure 4A, Table S4).** A comparative analysis shows that 19.4% of the total DE proteins identified in the cellular extract fraction fall into the insoluble category **(Figure 4B),** including proteins belonging to the visual cycle (RLBP1, DHRS3), protein folding (HSPB1) and splicing (PRPF31). Furthermore, GOBP enrichment analysis performed by Metascape showed that 934 DE proteins were involved in several key pathways including mRNA splicing, protein folding, response to ER stress and UPR **(Figure 4C),** some of which we showed earlier to be affected in RP11-RPE cells. Notably, several key spliceosomal tri-snRNP proteins including PRPF8, SNRNP200 (Brr2), PRPF4 and PRPF6, implicated in RP, as well as EFTUD2 (Snu114), USP39, SART1 and SART3 were amongst the splicing factors, whose amounts were significantly increased in the insoluble fraction **(Figure 4A, Table S4).** Moreover, the abundance of several HSPs is notably increased including components of the HSP90/R2TP chaperone system (HSP90, RUVBL1 and 2), previously implicated in the assembly of U4 and U5 snRNPs (38). Together these data suggest that key components of mRNA splicing, waste disposal, ER stress/UPR are deposited within the insoluble aggregates, preventing proper functions of these vital processes in RP11-RPE cells.

**Figure 4:**
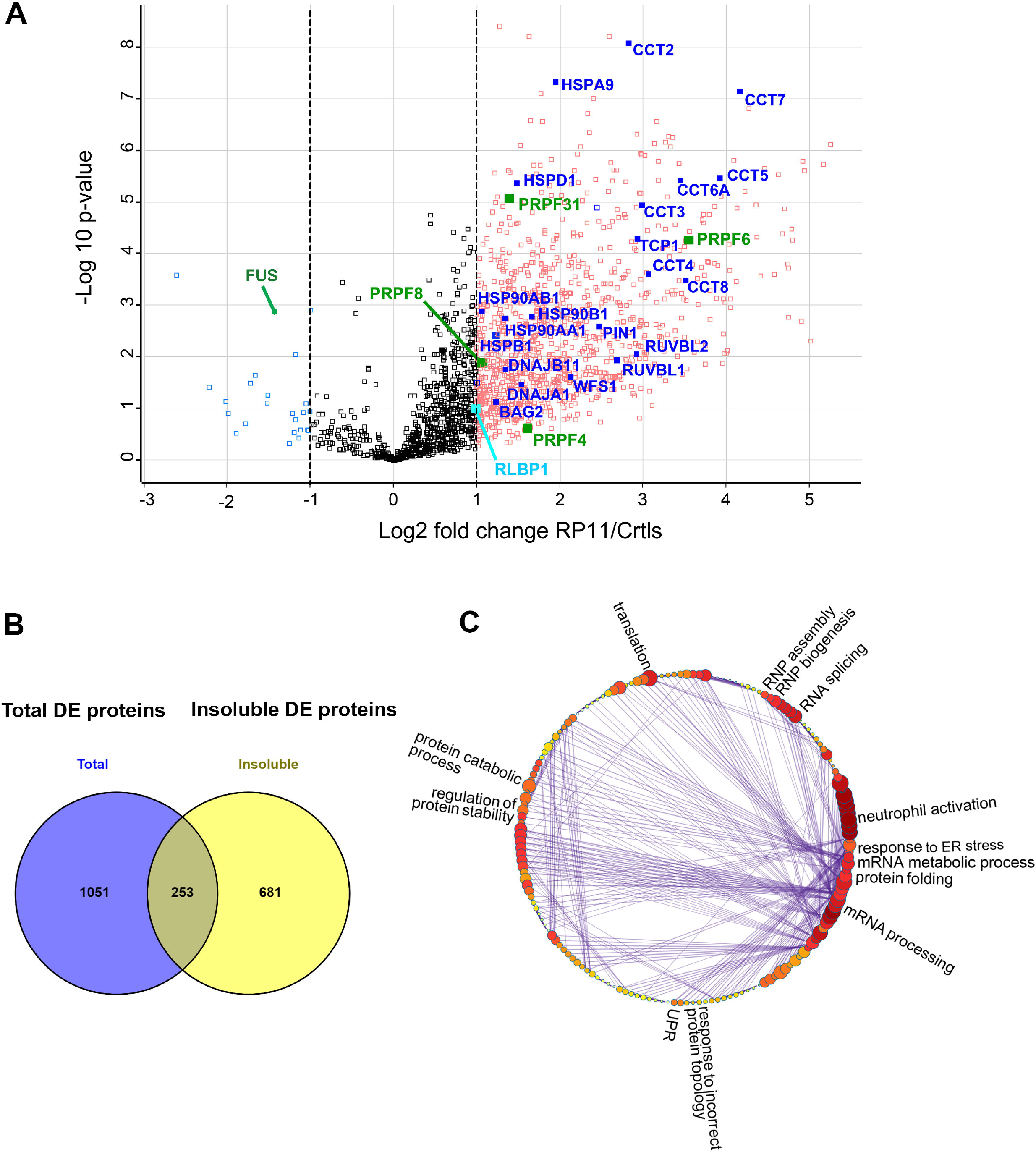
Differential protein abundance in the insoluble fractions of RP11-RPE cells. **A)** Volcano plot indicating the DE proteins in the insoluble fractions of RP11-RPE cells highlighting visual cycle (light blue), unfolded protein response (blue), visual perception (cyan) and spliceosome related (green) proteins. B) Venn diagram showing overlapping proteins between DE proteins detected from total cell extract (blue) and insoluble fractions (yellow) of RP11-RPE cells. **C)** Cluster illustrating the affected pathways of enriched GOBP insoluble proteins including RNA splicing, mRNA processing, mRNA metabolic process, protein folding, UPR, regulation of protein stability, protein catabolic process, translation and RNP assembly and biogenesis. A Log2 fold change cut off 1 was applied for the identification of significantly regulated proteins between insoluble fractions of control and RP11-RPE cells shown in Table S4.

### Accumulation of misfolded protein in RP11-RPE cells

Misfolded proteins which are unable to regain their normal conformation, are degraded by the ubiquitin-26S proteasome system (39). For the recognition of misfolded proteins by the 26S proteasome, misfolded protein need to be tagged by ubiquitin (mono-ubiquitination) or a chain of ubiquitin polypeptides (poly-ubiquitination) (40). Given that the GO analysis showed the enrichment of proteins involved in protein folding and proteolysis in RP11-RPE cells, we used the FK1 antibody that detects ubiquitin molecules, to identify ubiquitin-conjugated misfolded proteins. Western Blot analysis show an increased abundance of ubiquitin-conjugated proteins in RP11-RPE compared to the control cells **(Figure 5A).** The Western blot results were further validated by immunofluorescence analysis, which revealed an increased expression of ubiquitin-conjugated proteins in an aggregate-like pattern in patient specific RP11-RPE cells, but not in control cells **(Figure 5B).** These FK1 positive large aggregates were found in areas with disrupted tight junctions (ZO-1 staining) and devoid of nuclei. Furthermore, PRPF31 itself was associated with these FK1 positive aggregates **(Figure 5B).** These results suggest that misfolded protein destined for degradation are accumulated in the form of aggregates in RP11-RPE but not in control RPE cells, which suggests some dysfunction in the proteasome mediated degradation system in patient specific RP11-RPE cells.

**Figure 5.**
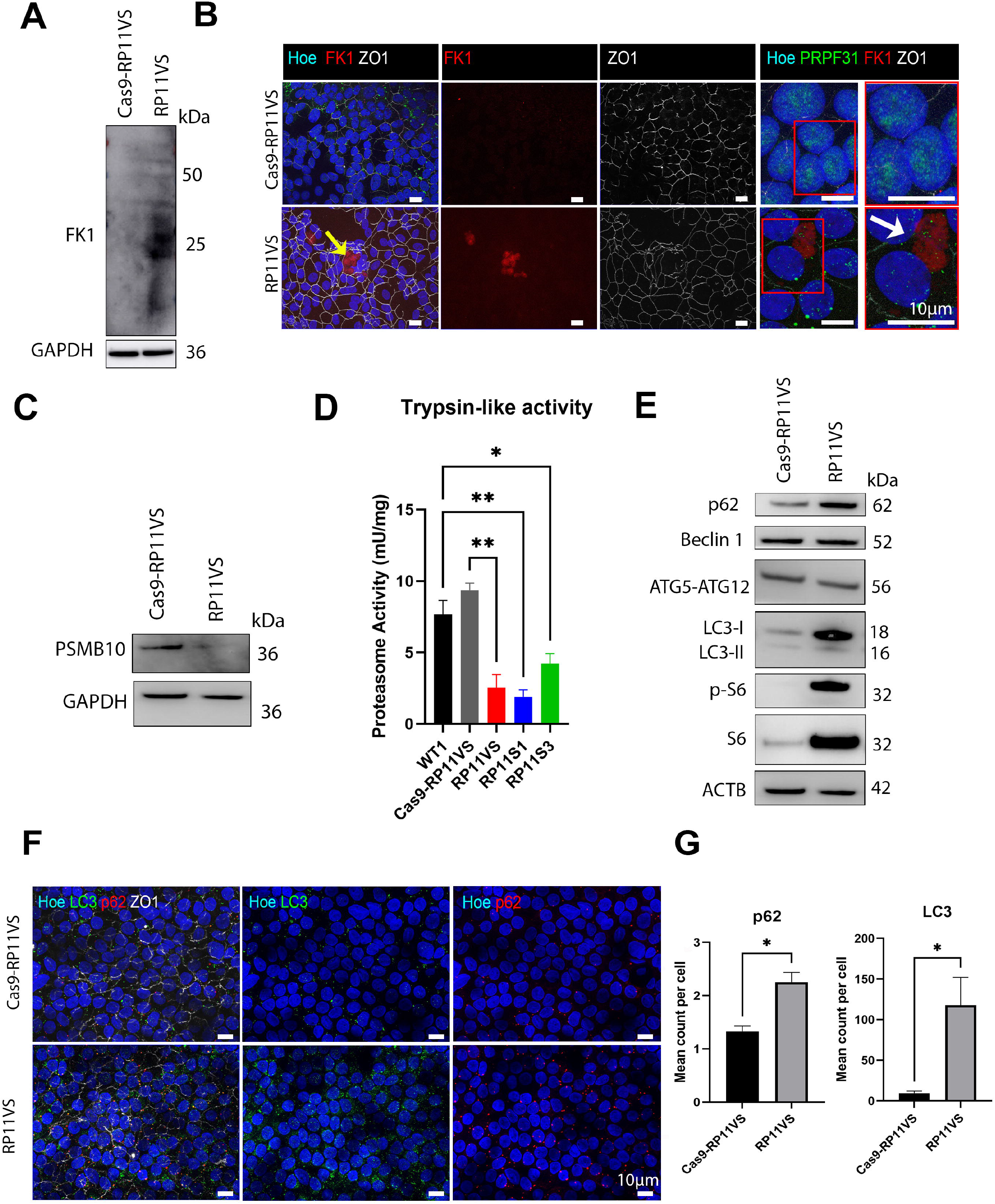
Accumulation of ubiquitin-conjugated proteins and dysfunction of the waste disposal mechanisms in RP11-RPE cells. **A)** Western blot showing upregulation of FK1 (ubiquitin-conjugated proteins) in RP11VS-RPE cells compared to control Cas9-RP11VS-RPE cells. GAPDH was used as a loading control. B) Immunostaining of RP11VS-RPE and control Cas9-RP11VS-RPE with FK1 (red) and ZO1 (white) showing accumulation of FK1 in RP11VS-RPE cells in the form of aggregates (shown by yellow arrow) but not in control cells. Magnified images show association of FK1 with PRPF31 (shown by white arrow). Cell nuclei were counterstained with Hoechst. Scale bars: 10 μm. Data obtained from RPE cells at week 12 of differentiation. C) Western blot of RPE samples showing downregulation of PSMB10 protein in RP11VS RPE cells compared to Cas9-RP11VS RPE cells. GAPDH was used as a loading control. D) Reduced Proteasome Trypsin-like activity in RP11-RPE cells compared to control RPE cells. Statistically significant differences are determined by t-test (*P<0.01, **P<0.001). Data are presented as mean ± SD, n=3. E) Western blot of key autophagic components showing upregulation of p62, LC3-I, p-S6 and S6 expression in RP11-RPE cells compared to Cas9-RP11VS RPE cells. Actin B was used as a loading control. F) Immunofluorescence analysis showed upregulation of p62 (red) and LC3 (green) in RP11-RPE compared to Cas9-RP11VS RPE cells. ZO1 (white) was used to define the tight junctions of RPE cells. Cell nuclei were counterstained with Hoechst. Scale bars: 10 μm. G) Quantification analysis of LC3 and p62. Statistically significant differences are indicated by t-test (*p< 0.01). Data are presented as mean ± SD, n=5.

To validate the presence of cytoplasmic aggregates in RP11-RPE cells, TEM analysis was performed revealing striking differences between control and RP11-RPE cells **(Figure S4).** Control RPE cells were characterised by normal cuboidal morphology, with no visible signs of degeneration. However, RP11-RPE cells were characterised by the presence of multivesicular bodies (MVBs) (indicated by red arrowheads) as well as big vacuoles filled with electrodense material suggesting the accumulation of cytoplasmic aggregates. Additionally, big gaps between RP11-RPE cells filled with debris (black arrowheads) were detected (denoted by red dotted lines) suggesting the accumulation of aggregates between RP11-RPE cells and disruption of tight junctions. Additionally, TEM images revealed the presence of expanded ER (red arrows), and the presence of stress vacuoles in patient - derived RP11 RPE cells **(Figure S4).**

### Dysfunction of the waste disposal mechanisms in RP11-RPE cells

In all tissues, the initial cellular defence mechanism is the ubiquitin proteasomal degradation pathway which together with the chaperone system degrades the majority of misfolded proteins (14). Degradation of the proteins occurs in the central 20S catalytic chamber which is composed of numerous subunits, although proteolytic activities are specifically performed by β1, β2, β5 subunits, which have caspase-like, trypsin-like, and chymotrypsin-like specificities (41). Our quantitative proteomic analysis revealed the downregulation of proteins responsible for the enzymatic activities of chymotrypsin-like (PSMB8), caspase-like (PSMB9) and trypsin-like (PSMB10) in RP11-RPE cells compared to the control cells **(Table S2).** To validate these results, Western blot was performed using an antibody to the PSMB10 protein (involved in trypsin-like activity), which was the most downregulated proteasomal related protein (mean LFC = −0.656). The results confirmed the downregulation of PSMB10 protein in RP11-RPE compared to control cells **(Figure 5C).** Thereafter, the trypsin-like activity was further evaluated showing a significant decrease in the activity in RP11-RPE compared to control cells **(Figure 5D).** Together these results suggest a significant downregulation of the proteosome proteolytic activity in RP11-RPE cells, which may impair the clearance of misfolded proteins, leading to their accumulation in the cytoplasm of patient-specific RPE cells.

When UPR and proteasome mediated degradation malfunction, activation of autophagy is observed (42). To this end, we assessed the expression of key components involved at different steps (autophagosome initiation, formation, and degradation) in the autophagy pathway. Our data show a significant increase in pS6, S6, p62, and LC3-I expression in RP11-RPE compared to control cells **(Figure 5E),** but no differences in expression of Beclin 1 or ATG5-ATG12, suggesting a block in the late stages of the autophagy-lysosome pathway. Immunofluorescence assays fully corroborated these data revealing increased p62 and LC3 expression in RP11-RPE cells **(Figure 5F, G).** These findings suggest the activation of the mTOR pathway and inhibition of autophagy, which together with dysfunction of the proteasome mediated degradation may lead to accumulation of misfolded proteins in cytoplasmic-like aggregates.

### Progressive accumulation of cytoplasmic aggregates in RP11-RPE cells

The accumulation of aggregate-prone proteins in the intracellular space has been associated with neurodegenerative diseases such as Alzheimer’s, Huntington’s and Parkinson’s, and Amyotrophic Lateral Sclerosis (ALS) (43) (44). All these diseases are characterised by a progressive pathology leading to the death of neurons. To investigate whether this also occurs in RP11 and assess the kinetics of aggregate accumulation in RP11-RPE cells, we performed immunofluorescence analysis at week 4, 8, and 12 post-plating of RPE cells in trans-wells, using antibodies to RLBP1, HSPB1, and FK1, which we showed earlier to accumulate in an aggregate-like pattern. Week 4 RP11-RPE cells displayed increased expression of RLBP1 and HSPB1 but not FK1 compared to control cells **(Figure S5).** Further increases in the expression of RLBP1, FK1 and HSPB1 were noticeable in RP11-RPE cells assessed at week 8 **(Figure S5).** RLBP1 was the only protein that showed accumulation in an aggregate-like pattern at this time point. At 12 weeks, all three markers displayed this aggregatelike accumulation but only in patient RP11-RPE cells, corroborating our earlier data **(Figure S5).** These results suggest a progressive accumulation of visual cycle proteins, chaperones, and misfolded-ubiquitinated proteins in the cytoplasm of RP11-RPE cells.

### Daily feeding of RP11-RPE cells with POSs accelerates cytoplasmic aggregate accumulation

One of the main functions of RPE cells is the engulfment and phagocytosis of POSs, which are shed from photoreceptors daily at a very high rate. To assess the aggregate accumulation under physiologically relevant conditions, the control and RP11-RPE cells differentiated in trans-wells for 4 weeks were fed with unlabeled POSS, and collected at 0, 48, 96 and 144 hours post feeding. Thereafter, the presence of intracellular aggregates containing RLBP1, HSPB1 and/or FK1 was analysed by immunofluorescence microscopy **(Figure S6A).** RP11-RPE cells fed with POSs at 0 hours, did not show any differences in RLBP1, HSPB1 or FK1 expression compared to control RPE cells. However, at 48 hours post feeding with POSs, RP11-RPE cells showed a significant increase in the accumulation of cytoplasmic aggregates containing RLBP1, HSPB1 or FK1 **(Figure S6A).** A further accumulation of proteins was observed at 96 and 144 hours post feeding, with RLBP1, HSPB1 and FK1 expressed as large aggregates in the cytoplasm of RPE cells disrupting the tight junctions between the cells **(Figure S6).** Additionally, the expression of LC3 and p62 was assessed, showing a gradual increase in the expression with time post POSs feeding **(Figure S6A).** However, in control RPE cells, no differences were observed in the expression of RLBP1, HSPB1 or FK1 at 0-, 48-, 96- and 144-hours post-feeding with POSs, suggesting that accumulation of cytoplasmic aggregates containing RLBP1, HSPB1 or FK1 is prevented in control cells **(Figure S6B).** Under normal steady state conditions, cytoplasmic aggregates should be digested by the autophagy or proteasome mediated degradation; however, the accumulation of p62 and LC3 observed only in RP11-RPE cells, suggests that the waste disposal mechanism is impaired or overwhelmed, resulting in aggregate accumulation. Altogether these results suggest that daily feeding of RP11-RPE cells with POSs accelerates the accumulation of cytoplasmic aggregates containing amongst others RLBP1, FK1 and HSPB1 proteins in RP11-RPE cells.

To assess the impact of cytoplasmic aggregate accumulation on cell survival, control and RP11-RPE cells were immunostained with a Caspase-3 antibody. Caspase-3 is synthesised as inactive proenzyme and is localised in the cytoplasm. However, during apoptosis, caspase-3 is proteolytically cleaved, activated, and translocated into the nucleus causing fragmentation of DNA and damaging essential cellular proteins including enzymes involved in DNA repair (45). Our data show that Caspase-3 is predominantly expressed in the cytoplasm of control RPE in contrast to RP11-RPE cells, where Caspase-3 is mainly expressed in the nucleus **(Figure S7A),** suggesting the activation of Caspase-3 in the latter.

Collectively our findings reveal the accumulation of cytoplasmic cellular aggregates containing amongst others both the wild type and mutated PRPF31 isoforms, misfolded ubiquitin-conjugated, chaperones and visual cycle proteins in RP11-RPE cells. The proteasome-mediated and autophagy degradation pathways are both impaired, resulting in the progressive accumulation of these cytoplasmic aggregates with time. This process is exacerbated under physiological conditions, leading to disruption of tight junctions and activation of cell death through apoptosis.

### Elimination of aggregates in RP11-RPE cells by pharmacological drugs

To assess whether cytoplasmic aggregate accumulation could be eliminated, we tested the following pharmacological drugs:

1. Arimoclomol, an investigational drug currently in phase III of clinical trial for ALS (46). This is a heat shock protein co-inducer that has been shown to enhance HSPs expression both *in vitro* (47) and *in vivo* (48) and to alleviate protein aggregation.
2. STF-083010 is a pharmacological compound that has been shown to decrease cell death and attenuate oxidative stress by targeting IRE1, which mediates the unfolded protein response (49). Salubrinal is another pharmacological drug that has been shown to have neuroprotective effects in ALS animal models (50, 51). Salubrinal protects the cells against apoptosis by blocking the PERK pathway, leading to further inhibition of the protein synthesis (52).
3. Rapamycin is a widely used pharmacological compound able to activate autophagy by inhibiting mTORC1 (53) (54). Trehalose, an AMPK-dependent activator (55) (56), has also been shown to activate autophagy and confer beneficial effects for the elimination of cytoplasmic aggregate-prone proteins. All these three compounds were used alongside Arimoclomol, STF-083010 and Salubrinal and vehicle controls (DMSO or water).

Their effects on cytoplasmic aggregates were assessed following 7-day treatment of RP11-RPE cells differentiated on trans-wells for 12 weeks. Immunostaining with RLBP1, HSPB1 and FK1 was performed to detect cytoplasmic aggregates. The data showed no significant difference in the accumulation of cytoplasmic aggregates containing RLBP1, HSPB1, FK1 or PRPF31 in RP11-RPE cells treated with Salubrinal, Arimoclomol, STF-083010 or trehalose **(Figure 6A).** Interestingly, a significant reduction in cytoplasmic aggregate accumulation was observed in RP11-RPE cells treated with Rapamycin only **(Figure 6A).** The same results were obtained in RP11-RPE cells from a second patients, harbouring a different *PRPF31* mutation **(Figure S8).** Also, western blot analysis was performed to assess whether autophagy was indeed activated in RP11-RPE cells after treatment with Rapamycin, demonstrating a decrease in the expression of pS6 and S6 in the rapamycin treated RP11-RPE cells, and an increase in the expression of LC3-I to LC3-II. However, no differences were detected in the expression of LAMP1 and p62. Furthermore, a decrease in the expression of RLBP1 and HSPB1 was detected after treatment of RP11S3 RPE cells with Rapamycin suggesting that Rapamycin induced the activation of autophagy in RP11-RPE cells leading to a significant elimination of aggregates containing RLBP1, HSPs or FK1 in RP11-RPE cells **(Figure 6B).**

**Figure 6:**
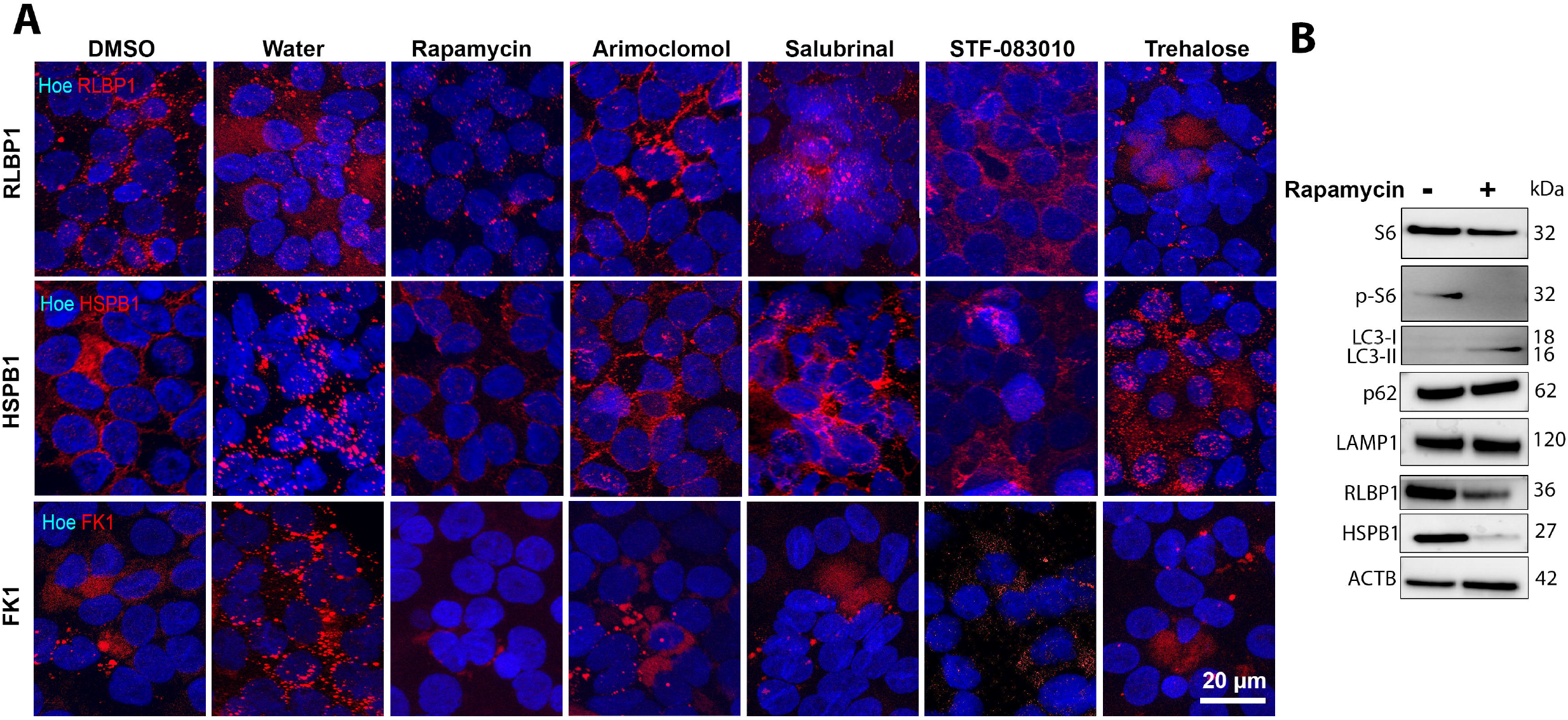
Elimination of aggregates in RP11-RPE cells through application of Rapamycin. Immunostaining of RP11S1-RPE cells showing a significant decrease of cytoplasmic aggregates containing RLBP1 (red), HSPB1(red) and FK1 (red) upon daily treatment with Rapamycin (500 ⊓M) for 7 days. Seven days treatment with Arimoclomol (1 μM), Salubrinal (25 μM), STF-083010 (50 μM) and trehalose (50 mM) showing no significant differences. DMSO or distilled water were used as vehicle controls. Cell nuclei were counterstained with Hoechst. Scale bars: 20 μm. **B)** Western blot of 7 days Rapamycin treated vehicle treated RP11S1-RPE cells showing the decrease in the expression of S6, p-S6 RLBP1, HSPB1, and activation of LC3-II in RP11S3-RPE cells.

Thereafter, we assessed the effects of Rapamycin treated RP11-RPE cells on cell survival after 7 days treatment of RP11-RPE cells with vehicle or Rapamycin respectively. In rapamycin treated RP11-RPE cells we observed reduced nuclear localisation of Caspase-3, indicating a positive effect in cell survival **(Figure S7B).** These results suggest that Rapamycin has a positive impact in the elimination of cytoplasmic aggregates, potentially leading to enhanced cell survival.

### Differential expression and aggregate formation of key proteins implicated in Retinitis Pigmentosa

We next searched in our proteomics datasets for other retinal-specific proteins DE in PP11-RPE or retinal organoids or deposited within the aggregates. To this end, Metascape was used to retrieve DE genes associated with diseases from the DisGeNET database **(Table S2, S3, S4).** This analysis revealed RP as one of the enriched disorders in both datasets of RPE and retinal organoids DE proteins.

In RPE, besides RLBP1, we found also upregulation of Bestrophin-1 (BEST1), an anion channel expressed in RPE and essential for calcium signaling, whose mutations lead to RP (51). PROS1 levels that has been linked to juvenile RP is increased within the insoluble RPE fraction (57). Retinol dehydrogenase 5 (RDH5) previously linked to RP was significantly upregulated in the total and insoluble RPE fractions (58). *PHYH* whose mutations cause adult Refsum disease, a peroxisomal disorder with numerous features including RP, is highly upregulated in the total cellular and insoluble fractions. In addition, differential expression of key RPE proteins RGR, RPE65 and MYO7A was noticed in our RPE proteomics dataset.

In retinal organoids we detected upregulation of HGSNAT, a lysosomal acetyltransferase whose variants are the cause of RP-73, and HKDC1, a hexokinase localized to the photoreceptor inner segment and associated with autosomal recessive RP, as well as Retinol dehydrogenase 11 (RDH11) associated with syndromic RP, and the MYO7A protein involved in Usher syndrome with an RP phenotype. GNAT2, a gene associated with cone-rod dystrophies is the most significantly downregulated protein essential for the phototransduction pathway in RP11 retinal organoids (59). Together our data demonstrate dysregulation of expression and/or localisation and/or aggregation of multiple RP genes in both RP11-RPE cells and retinal organoids ensuing from *PRPF31* mutations.

## Discussion

In previous work, using an iPSC-disease modelling approach, we have shown that mutations in the spliceosomal U4/U6 snRNP-specific protein PRPF31, result in global splicing changes specifically in retinal cells and RPE (10). To fully understand the impact of such global splicing dysregulation on the patient derived RPE and retinal cells, we have undertaken a detailed quantitative proteomic and immunofluorescence analysis, as well as biochemical analysis of these cells harbouring *PRPF31* mutations. These new findings have shown the predominant accumulation of the mutant PRPF31 protein as aggregates in the cytoplasm of RPE cells and its significant presence in the cytoplasm of retinal cells. This results in reduced nuclear levels of wild type isoform in RP11 patient RPE and retinal cells, leading to reduced amounts of U4/U6 snRNPs and accumulation of U5 in Cajal bodies, thus defects in tri-snRNP formation as well as smaller size of nuclear speckles. These assembly defects also result in reduced amounts of active spliceosomes in both RPE cells and retinal organoids. Quantitative proteomics revealed that several key cellular processes in addition to mRNA splicing, namely waste disposal and unfolded protein response were significantly affected resulting in progressive accumulation of insoluble aggregates containing key components of each of the pathways within RPE cells, that affected cell viability. Aggregate formation also affected the expression and/or localization of key proteins of the retinoid metabolism and visual perception pathways including RLBP, which accumulated in the PRPF31 cytoplasmic aggregates. Furthermore, proteomic analysis of insoluble aggregates in RP11-RPE cells identified the presence of other core components of the tri-snRNP including PRPF6, PRPF4 and PRPF8 (all of which are linked to adRP) together with chaperone proteins important for U4/U6 and U5 snRNPs assembly. Strikingly, activating autophagy via application of Rapamycin, resulted in reduction of these cytoplasmic aggregates and correct nuclear localization of PRPF31 in RP11-RPE cells that improved cell survival **(Figure 7).**

**Figure 7:**
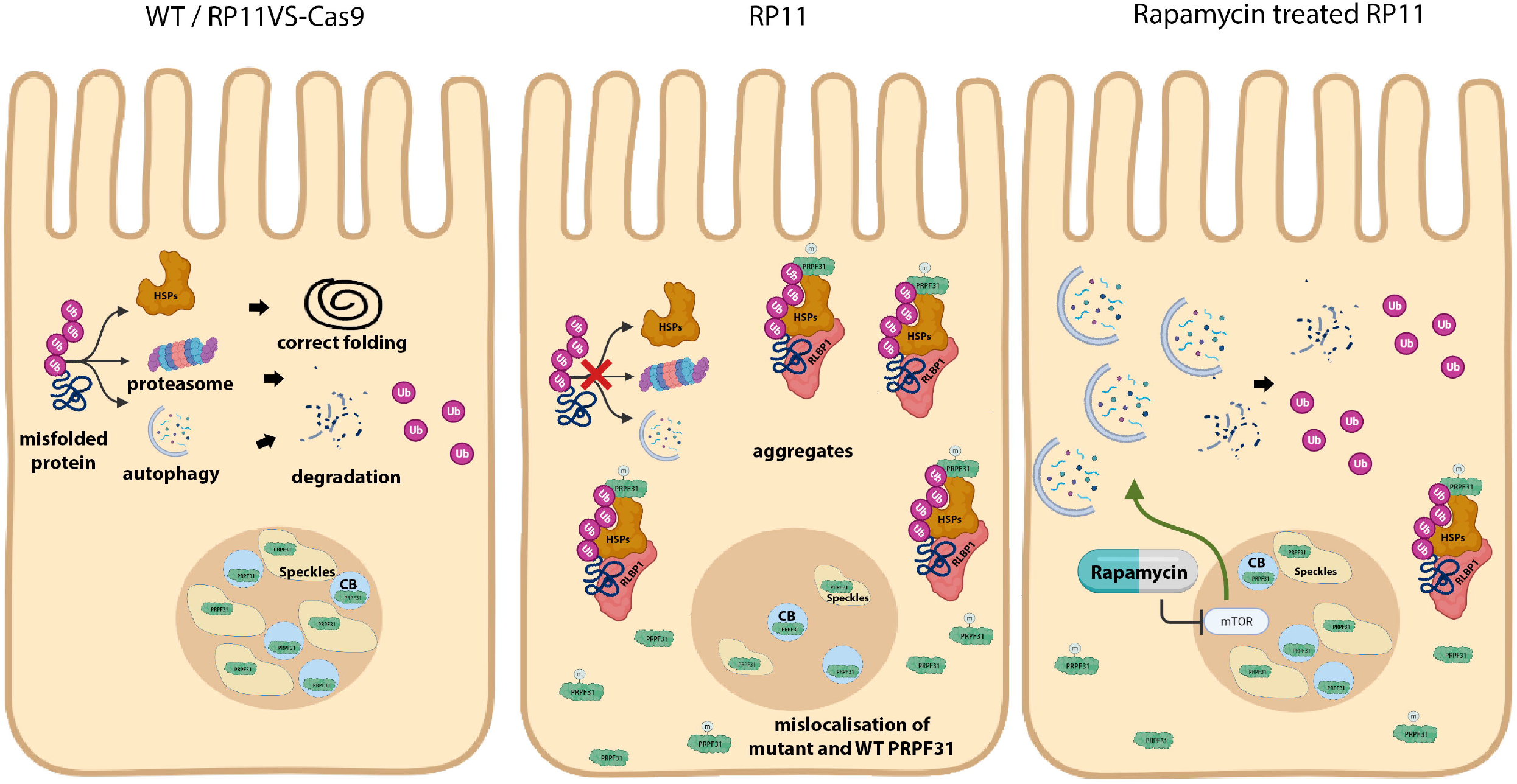
Schematic presentation showing localisation of mutant PRPF31 in the cytoplasm of RP11-RPE cells and accumulation of aggregates containing HSPs, visual cycle and ubiquitin conjugated proteins, which were much reduced upon application of autophagy activator, Rapamycin. m - mutant PRPF31, CB-Cajal Bodies, Ub-ubiquitin.

Aggregation of PRPFs in retinal photoreceptors and RPE cells has been reported previously and shown to be cell type specific. For example, the mutant PRPF3^T494M^ amasses in big aggregates in the nucleolus region (11), causing mislocalisation of splicing factors that may be detrimental for photoreceptor cells. In contrast, the mutant PRPF31 protein forms insoluble aggregates in the cytoplasm of RPE cells of Prpf31^A216P/+^ mice (13), decreasing the protein levels of this splicing factor in the nucleus. Our data show reduced nuclear expression of the wild type PRPF31 and cytoplasmic localisation of mutant PRPF31 protein, corroborating the mouse RPE cell studies. Knockdown studies in HeLa cells have demonstrated that lack of Prpf31 or Prpf6 leads to accumulation of U4/U6 di-snRNPs in Cajal bodies and inhibition of tri-snRNP formation (16). Nonetheless, a detailed analysis of the impacts of *PRPF* mutations on tri-snRNPs formation within Cajal bodies of retinal cells, spliceosome assembly in nuclear speckles and formation of active spliceosomes directly on patient specific RPE and retinal cells has not been reported previously. Using a combination of RNA-FISH and immunofluorescence microscopy, we demonstrate a significant defect in the tri-snRNP assembly in patient photoreceptor cells leading to a decrease in U4/U6 di-snRNPs and accumulation of U5 snRNP in Cajal bodies, the nuclear membrane-less organelles where tri-snRNPs are assembled and fully matured (15). Strikingly, glycerol gradient fractionation combined with Northern and Western blotting, demonstrated for the first time that the mutant PRPF31 is not incorporated into the spliceosomal complexes in RP11-RPE cells. Moreover, an accumulation of U4 and U6 and slow sedimenting PRPF31 in the gradient top fractions are observed in the RP11 patient RPE cells. These defects in tri-snRNP formation in RP11 cells most likely will lead to perturbed pre-mRNA splicing. Consistently, RP11 retinal organoids and RPE cells showed altered morphology of nuclear speckles, the nuclear compartments for storage of splicing factors and associated with post-transcriptional splicing, as well as reduced localisation of PRPF31 to these compartments suggesting a reduction in active spliceosomes. To prove this, we immunostained the retinal organoids and RPE cells with an antibody raised against a phosphorylated form of the U2-specific protein SF3B1, which is phosphorylated in activated (B^act^) or catalytically active (complex C) spliceosomes. A significant reduction in staining was observed in RP11 retinal organoids, demonstrating reduced levels of active spliceosomes and splicing activity in patient cell lines. A mild reduction for p-SF3B1 staining in RP11-RPE cells was also observed, suggesting a reduced splicing activity in these cells compared to controls. Notably, gradient profiles showed lower levels of tri-snRNPs in RPE cells compared to retinal organoids, which could further imply that even mild changes in splicing activity would impact RPE cells more significantly than retinal cells. This however needs further investigations in RP11-RPE cells as well as other PRPF-RPE cells, which is currently ongoing in our groups.

In addition to splicing dysregulation, our proteomics data revealed an enrichment of proteins involved in the retinoid metabolic process and visual perception. Of particular interest in the retinoid metabolic and visual perception is the Retinaldehyde-binding protein (RLBP1), which is significantly overexpressed in RP11-RPE cells, and found in the form of cytoplasmic aggregates. RLBP1 is a retinoid-binding protein expressed in RPE and Müller glia cells and involved in the conversion of 11-*trans*-retinal to the light sensitive 11-*cis* retinal (60). Mutations in *RLBP1* cause a range of retinopathies including retinitis punctata albescens (RPA), Bothnia-type dystrophy (BD), Newfoundland rod-cone dystrophy (NFRCD), retinitis pigmentosa (RP) and fundus albipunctatus (FA) (61). Based on our data, our hypothesis is that accumulation of RLBP1 in the insoluble aggregates, reduces the available RLBP1 protein needed for recycling of retinal, impacting directly on the visual cycle and the phototransduction pathway.

Studies performed in the RPE of Prpf31^A216P/+^mice have revealed overexpression of heat shock protein 70 (Hsp70) family, and its co-localisation with mutant Prpf31 in the insoluble cytoplasmic aggregates. It is reasonable to assume that protein aggregation of mutant PRPF31 is responsible for activating the chaperone response. Corroborating these studies, our data also revealed significant changes in expression of several HSPs and overexpression of HSPA4L, HSPB1 and HSPA2, that accumulated in the form of aggregates in association with PRPF31 in in RP11-RPE cells. Remarkably, HSP aggregation between photoreceptor cells was also observed in RP11 retinal organoids. HSPs facilitate protein homeostasis and cell protection against damaged proteins or aggregation of harmful denatured proteins. Their upregulation in RP11-RPE cells could be a direct response of global splicing deregulation, which is bound to result in misfolded or aggregated proteins, as shown in our study. Previous studies have reported that under stress conditions (e.g. hyperglycemia) or upon mutations, soluble HSPs are depleted, becoming unavailable to target other proteins, affecting further their functionality (62) and leading to the accumulation of insoluble aggregates in the retina (62). Our data demonstrates preferential HSP accumulation in the insoluble aggregates of RP11-RPE cells, suggesting a change in their solubility and function, which could be due to “overwhelming” of HSP response by the large amount of misfolded or aggregates proteins resulting from the global spliceosome dysfunction.

The formation of cytoplasmic aggregates is a common hallmark event of many neurodegenerative diseases and is associated with protein misfolding (1). Usually, in response to misfolded protein, cellular defence mechanisms like molecular chaperons such as HSPs are activated. However, when quality control systems are incapable to restore the normal conformation of denature proteins and thus are overloaded with excessive amounts of denatured proteins, then it is possible that the solubility of HSPs can be transformed from a soluble state to an insoluble form as shown in our study. When HSPs fail to restore misfolded proteins, denatured or aggregated proteins are tagged with ubiquitin (Ub) and are directed to the Ub-proteasome system (UPS) for proteolytic degradation (43). Our results revealed increased protein ubiquitination and a reduced enzymatic activity (trypsin-like) of the proteasome degradation system in RP11-RPE compared to control RPE cells. The reduced proteasome activity revealed in our results is also a common characteristic feature of other NDs such as AD, PD, ALS, and HD (43). Particularly in AD disease, various studies have shown that tau aggregates can bind to the recognition site of the 19S catalytic particle of the proteasome inducing protein congestion that further leads to impairment of protein degradation (63). Also, misfolded prion proteins (β-sheet-rich PrP) in Prion disease disrupt the opening of the 20S proteasome particle, thus inhibiting the function of the 26S proteasome (64). This association of pathogenic protein aggregates and reduced or blocked proteasome activity has been reported by several studies focusing on the AD (65, 66). Our data mirror these findings, however, in the case of RP11-RPE cells, it is unknown whether the accumulation of misfolded/aggregated proteins is a result of decreased proteasome activity or whether the proteasome system is incapable of coping with the burden of misfolded, ubiquitinated and aggregated proteins resulting from global spliceosome dysregulation.

Under stress conditions when the chaperone and proteasome systems are overwhelmed, clearance of cytoplasmic aggregates is facilitated by autophagy, where cytoplasmic substrates are engulfed and degraded into amino acids (43). Although most proteins in the human genome can be successfully degraded from the cells, genetic defects can affect their conformation leading to the formation of aggregates. For example, mutations of huntingtin proteins inhibit the proteolytic machinery and induce accumulation of cytoplasmic aggregates in patients with HD (67). This is characterised by an increase of key autophagic components such as p62, LC3 in HD mouse models due to the impairment of cargos to be directed to autophagic vacuoles for degradation (68). Likewise, our results demonstrate an upregulation of p62 and LC3 expression in RP11-RPE cells compared to control RPE cells suggesting an impairment of autophagy. We hypothesise that this combined dysfunction of proteasome and autophagy mediated degradation together with accumulation of HSPS in the insoluble fractions leads to the accumulation and growth of larger aggregated proteins, which may be cytotoxic to the cells. This is a characteristic of many NDs such as HD, AD, PD, and ALS, where mutant aggregated proteins become resistant to degradation inducing cytotoxicity and neuronal cell death (43). To this end it has been reported that neuronal cells of AD patients (69) have increased levels of Caspase-3, associated with degeneration of synapses and a decrease in synaptic plasticity, suggesting that intracellular protein deposits might disrupt the normal function of neurons, inducing stress which in turn leads to the initiation of cell death. Similarly, our results have shown the progressive accumulation of large cytoplasmic aggregates containing the mutant PRPF31 itself as well as misfolded, ubiquitin conjugated, and visual cycle proteins are found in areas with disrupted tight junctions and increased Caspase 3 activity. Tight junction disruption has been reported by other studies associating RPE remodeling with retinal degeneration (70). Collectively these results suggest that continuous accumulation of aggregates in patient-specific RP11-RPE cells impairs RPE cell survival.

One potential strategy to enhance the degradation of misfolded proteins is to induce activation of autophagy. Many small molecules have been developed to activate autophagy and induce clearance of pathogenic proteins, and the most widely known is by inhibiting mTORC1 by Rapamycin. It has been shown that Rapamycin can effectively reduce cytoplasmic mutant proteins such as α-synuclein (71), huntingtin (72), or tau mutant proteins (73) from the brains of transgenic mouse models. Our studies show that cytoplasmic aggregates and cell death can indeed be reduced upon treatment of RP11-RPE cells for 7 days with Rapamycin. However, other pharmacological treatments targeting autophagy (trehalose), ER stress (Salubrinal and STF-083010) or inducers of HSPs such as Arimoclomol had no beneficial effects in reducing the volume of cytoplasmic aggregates in RP11-RPE cells. Our results are in agreement with other studies using Drosophila (74) and mouse models (75) associated with aggregate-prone proteins, showing the clearance of cytoplasmic aggregates after treatments with Rapamycin, demonstrating its potential therapeutic use in diseases associated with aggregate accumulation (74, 76). The improved aggregate clearance by activation of autophagy via Rapamycin administration indicates that it is the progressive aggregate accumulation that overburdens the waste disposal machinery rather than direct PRPF31 initiated misplicing. Hence therapeutic strategies aiming at waste disposal activation present an important approach that needs to be investigated in close conjunction with gene therapy studies.

## Acknowledgments

The authors are grateful for funding support from Retina UK (GR595, GR584), MRC UK (MR/T017503/1), Fight for Sight (1456/1457) and ERC (CoG_614620). We would like to thank Newcastle University Bioimaging facility for help with imaging assays. Acknowledgment is made to Bio Render (www.biorender.com). Figure 7 was created with BioRender.com.

## Author contributions

Maria Georgiou: Collection and/or assembly of data, data analysis and interpretation, figure preparation, manuscript writing

Chunbo Yang: Collection and/or assembly of data, data analysis

Robert Atkinson: Collection and/or assembly of data

Kuan-Ting Pan: Collection and/or assembly of data, data analysis

Adriana Buskin: Collection and/or assembly of data

Marina Moya Molina: Collection and/or assembly of data

Joseph Collin: Collection and/or assembly of data, fund raising

Jumana Al-Aama: Conception and design, fund raising

Franziska Goetler: data analysis

Sebastian E. J. Ludwig: Collection and/or assembly of data

Tracey Davey Collection and/or assembly of data

Reinhard Lührmann: Conception and design, fund raising

Sushma Nagaraja-Grellscheid: Conception and design, data analysis, fund raising

Colin Johnson: Conception and design, fund raising

Robin Ali: Conception and design, fund raising

Lyle Armstrong: Conception and design, fund raising

Viktor Korolchuk: data analysis and interpretation

Henning Urlaub: Conception and design, fund raising

Sina Mozaffari-Jovin: Conception and design, Collection and/or assembly of data, data analysis and interpretation, figure preparation, manuscript writing, fund raising

Majlinda Lako: Conception and design, data analysis and interpretation, manuscript writing data dissemination, overall project management and fund raising.

All authors read and approved the final manuscript.

## Disclosure

The authors declare no conflict of interest. The funders had no role in study design, data collection and analysis, decision to publish, or preparation of the manuscript.

## Supplementary Figures and Tables

**Figure S1. Localisation of mutant and WT PRPF31 in the cytoplasm of RP11-RPE cells.** RP11-RPE cells were immunostained with an anti-PRPF31 N terminus (green) and ZO1 (red) antibodies. Cell nuclei were stained with Hoechst. Immunofluorescence analysis showed localisation that PRPF31 protein is predominantly located in the cytoplasm in an aggregate like pattern in RP11VS and RP11S3 RPE cells. Scale bars: 20 μm.

**Figure S2: Analysis of snRNP levels in RP11VS and Cas9-RP11VS RPE cells and retinal organoids.** Northern blots of glycerol gradient fractions showing the level of snRNAs in Cas9-RP11VS and RP11VS RPE cells and retinal organoids. Total RNA was isolated from each sample and snRNA levels were analysed by denaturing PAGE followed by Northern blotting using radiolabelled probes against U1, U2, U4, U5 and U6 snRNAs and autoradiography (top).

**Figure S3: GO analysis reveals enrichment of proteins involved in RNA splicing, autophagy and lysosome, endoplasmic reticulum lumen and unfolded protein response and visual perception pathway in RP11-retinal organoids. A)** GO analysis showing DE proteins between control and RP11 retinal organoids involved in RNA splicing, highlighting with black circle PRPF31 protein. B) GO analysis showing DE proteins between control and RP11 retinal organoids involved in autophagy and lysosome. C) Western blot showing the downregulation of PRPF31 protein in RP11 retinal organoids compared to unaffected (WT1) and isogenic control organoids. Western blot showing upregulation of LAMP1 and LAMP2 in RP11 retinal organoids compared to unaffected (WT1) and isogenic control organoids. ACTB was used as a loading control. GO analysis identifies DE proteins between RP11-retinal and control organoids belonging to the D) Endoplasmic reticulum lumen, E) Unfolded protein response and F) visual perception.

**Figure S4. Ultrastructural analysis of RP11- and control RPE cells.** Representative TEM images of control and RP11-RPE cells showing the accumulation of amorphous electrodense material (black arrowheads) located between patient-derived RPE cells (red dotted line), presence of stress vacuoles (V), expanded endoplasmic reticulum (ER) (red arrow), and multivesicular bodies (MVBs) (red arrowhead). Control RPE cells were characterised by normal cuboidal morphology, with no signs of degeneration.

**Figure S5: Progressive accumulation of cytoplasmic aggregates in RP11-RPE cells.** Immunofluorescent images of control (RP11VS-Cas9) and RP11 (RP11VS) RPE cells at week 4, 8 and 12 showing a gradual increase and accumulation of RLBP1, HSPB1, and FK1 (red), forming large aggregates. Cell nuclei were counterstained with Hoechst. Scale bars: 20 μm.

**Figure S6. Evaluation of cytoplasmic aggregates in RP11 and control RPE cells after feeding with POSs.** Immunofluorescent images of RP11-RPE and Cas9-RP11-RPE cells which were fed daily with unlabelled POSs (20 POSs/cell) and collected at different time points (0, 48, 96 and 144 hours post feeding). Representative immunofluorescence images of RP11-RPE showing the accumulation of cytoplasmic aggregates (containing RLBP1, HSPB1 and FK1 (red) and autophagic markers, LC3 (green) and p62 (red) but no differences in the expression of RLBP1, HSPB1 and FK1 (red) or in the expression of LC3 and p62 were detected after feeding with POSs at 0, 48, 96 and 144 hours. Cell nuclei were counterstained with Hoechst. Scale bars: 20 μm.

**Figure S7. Activation of caspase-3 in RP11-RPE cells and the effects of Rapamycin in RP11VS-RPE cells. A)** Immunofluorescence images of RP11VS and RP11VS-Cas9 RPE cells stained with Caspase-3 (red) and ZO1 (white) showing nuclear localisation of caspase-3 in RP11-RPE cells (shown by white arrow) and cytoplasmic expression in control RPE cells. Magnified images show co-localisation of Caspase-3 with Hoechst in RP11VS-RPe cells. Cell nuclei were counterstained with Hoechst. Scale bars: 10 μm. Data obtained from RPE at week 12 of differentiation B) Immunostaining of Rapamycin and vehicle treated RP11VS-RPE cells with Caspase-3 (red) and ZO1 (white) showing reduced nuclear levels of cleaved Caspase-3 in the nucleus of rapamycin treated RP11VS-RPE cells. Magnified images show the expression level of Caspase-3 in vehicle and rapamycin treated RP11VS-RPE cells (shown by white arrow).

**Figure S8: Elimination of aggregates in RP11-RPE cells by pharmacological drugs in RP11S3-RPE cells.** Immunostaining of RP11S3-RPE cells showing a significant decrease of cytoplasmic aggregates containing RLBP1, HSPB1 and FK1 (red) upon daily treatment with Rapamycin (500 nM) for 7 days. No significant differences were detected after 7 days treatment with Arimoclomol (1 μM), Salubrinal (25 μM), STF-083010 (50 μM) and trehalose (50 mM). Cell nuclei were stained with Hoechst. Scale bars: 20 μm.

**Table S1: Summary of antibodies used in this study.**

**Table S2: DE proteins between control and RP11 RPE cells together with GOBP, GOCC and GO-DisGeNET analyses.**

**Table S3: DE proteins between control and RP11 retinal organoids together with GOBP, GOCC and GO-DisGeNET analyses.**

**Table S4: DE proteins between control and RP11-RPE cells found in the insoluble fraction.**

## Notes

### Competing Interest Statement

The authors have declared no competing interest.

