## Supplementary figures and images for "Progressive protein aggregation in PRPF31 patient retinal pigment epithelium cells: the mechanism and its reversal through activation of autophagy"

### Supplementary Figure 1

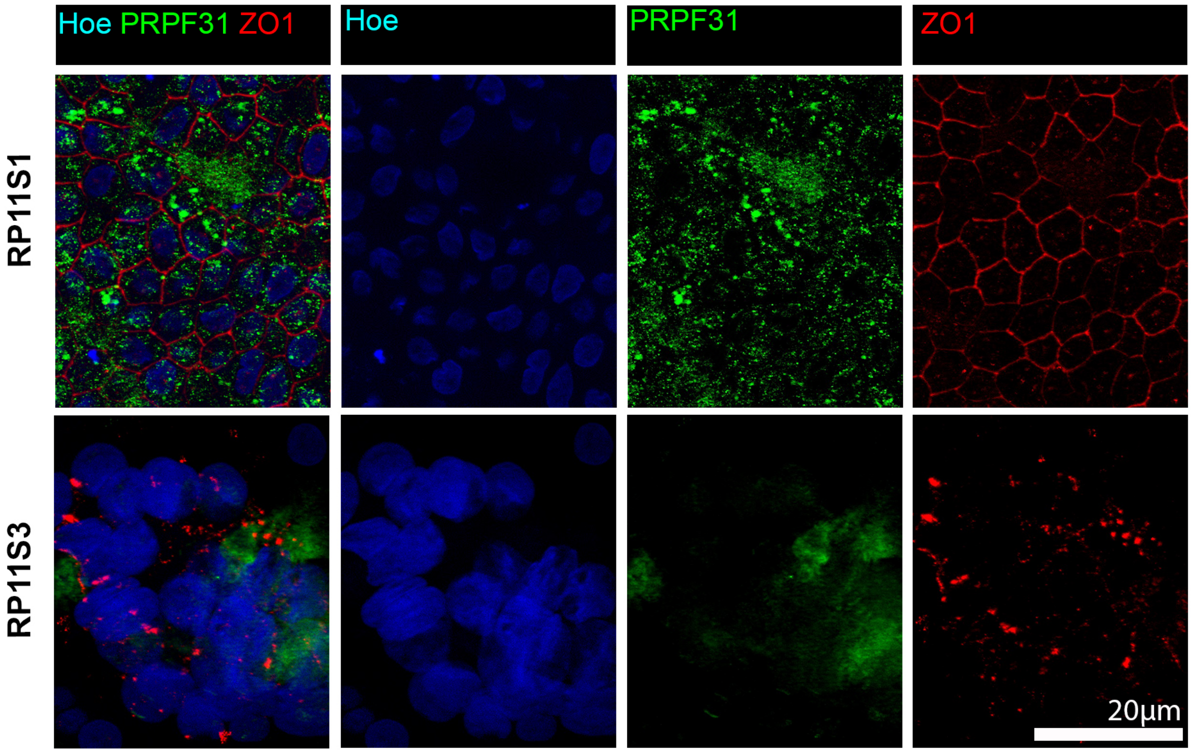


#### Figure S1

### Supplementary Figure 2

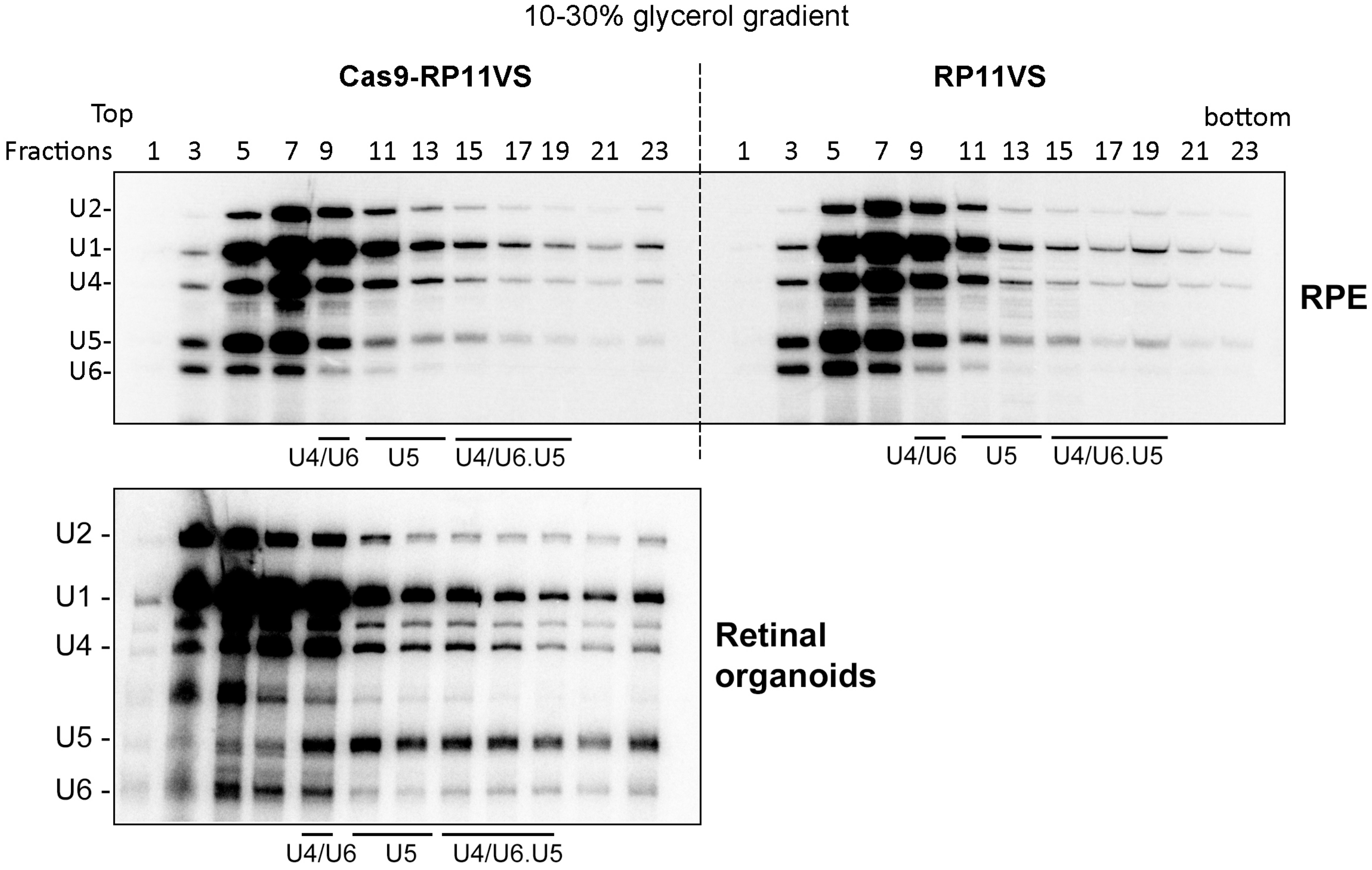

### Supplementary Figure 3

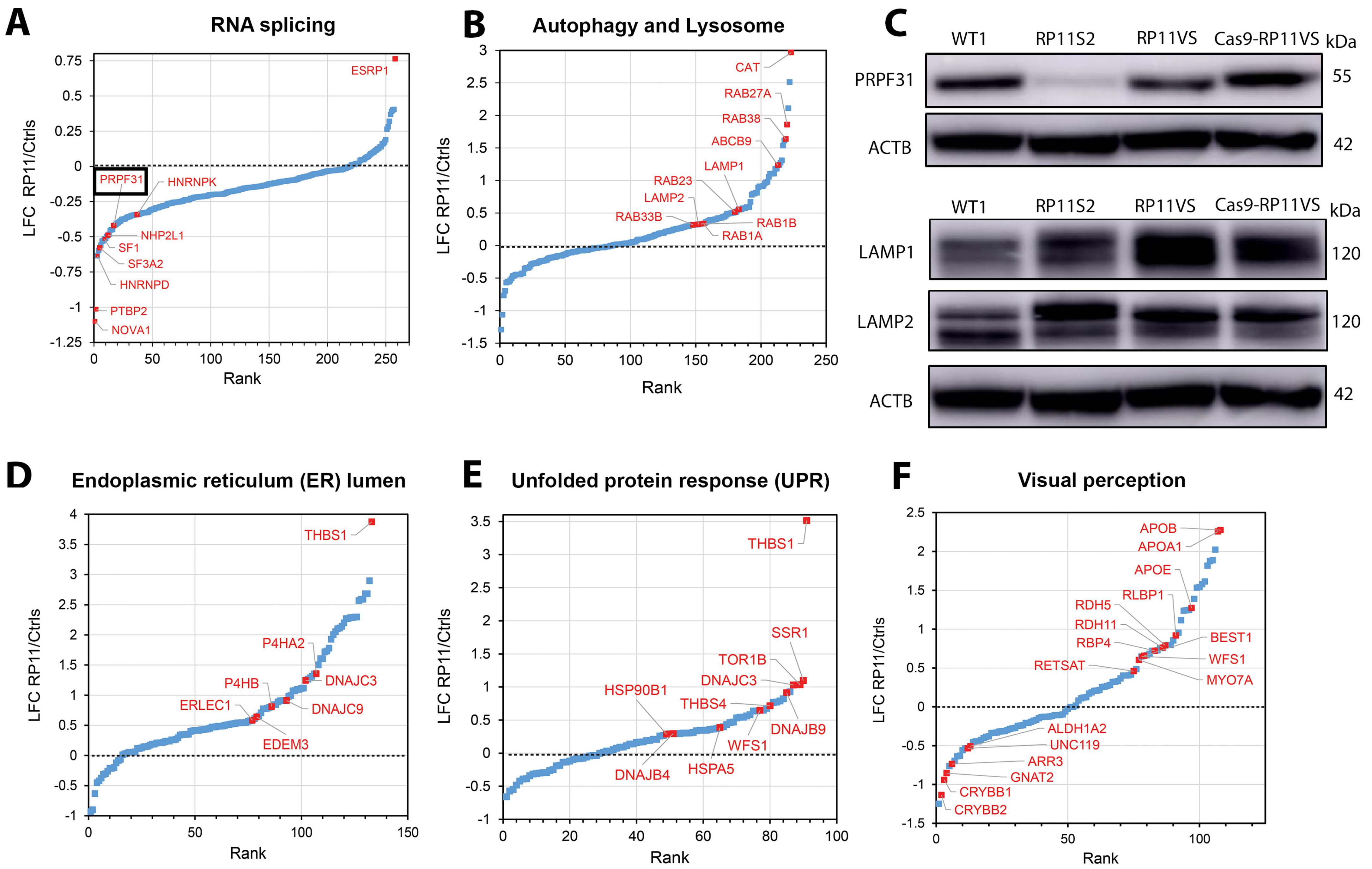

### Supplementary Figure 4

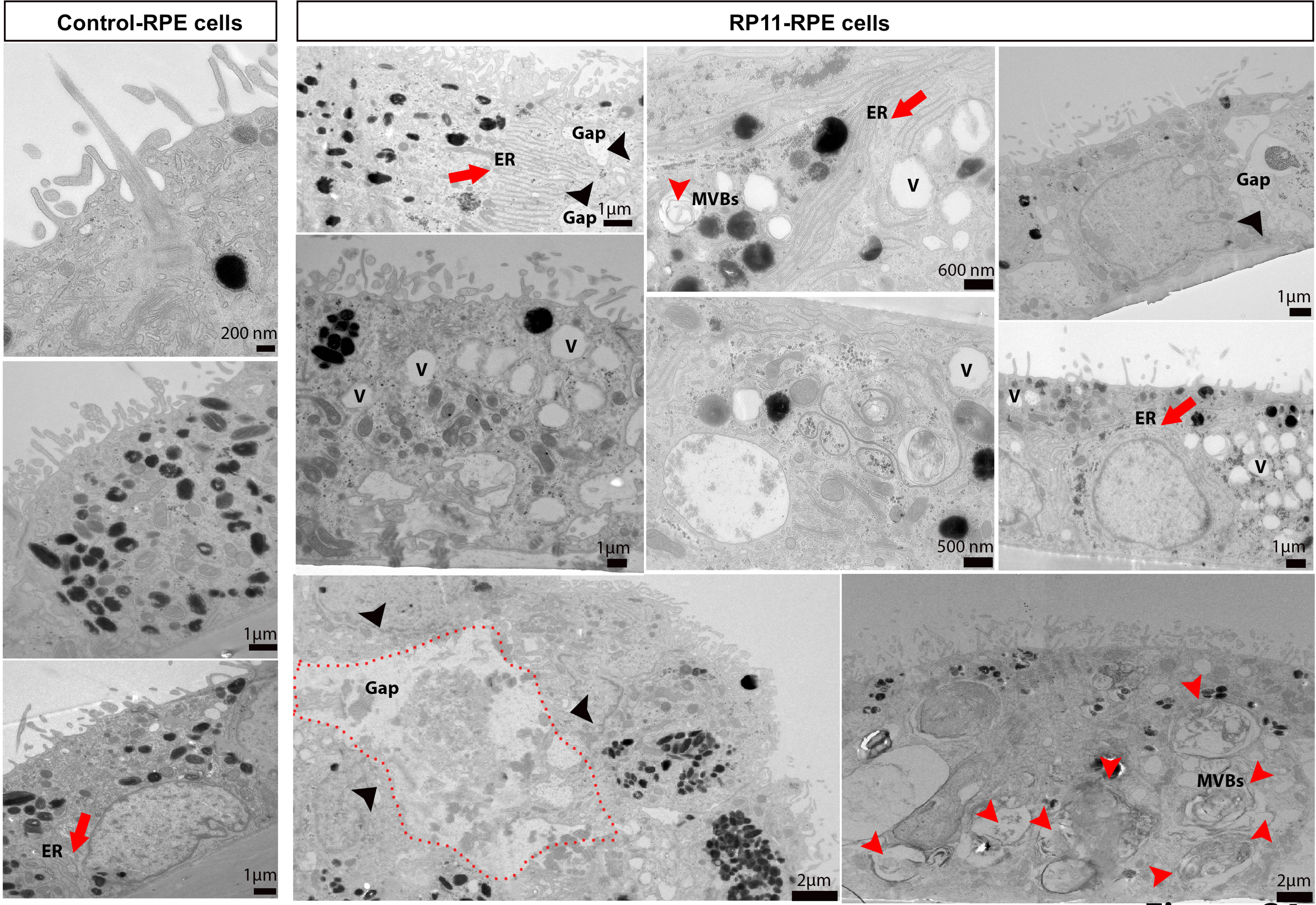

### Supplementary Figure 5

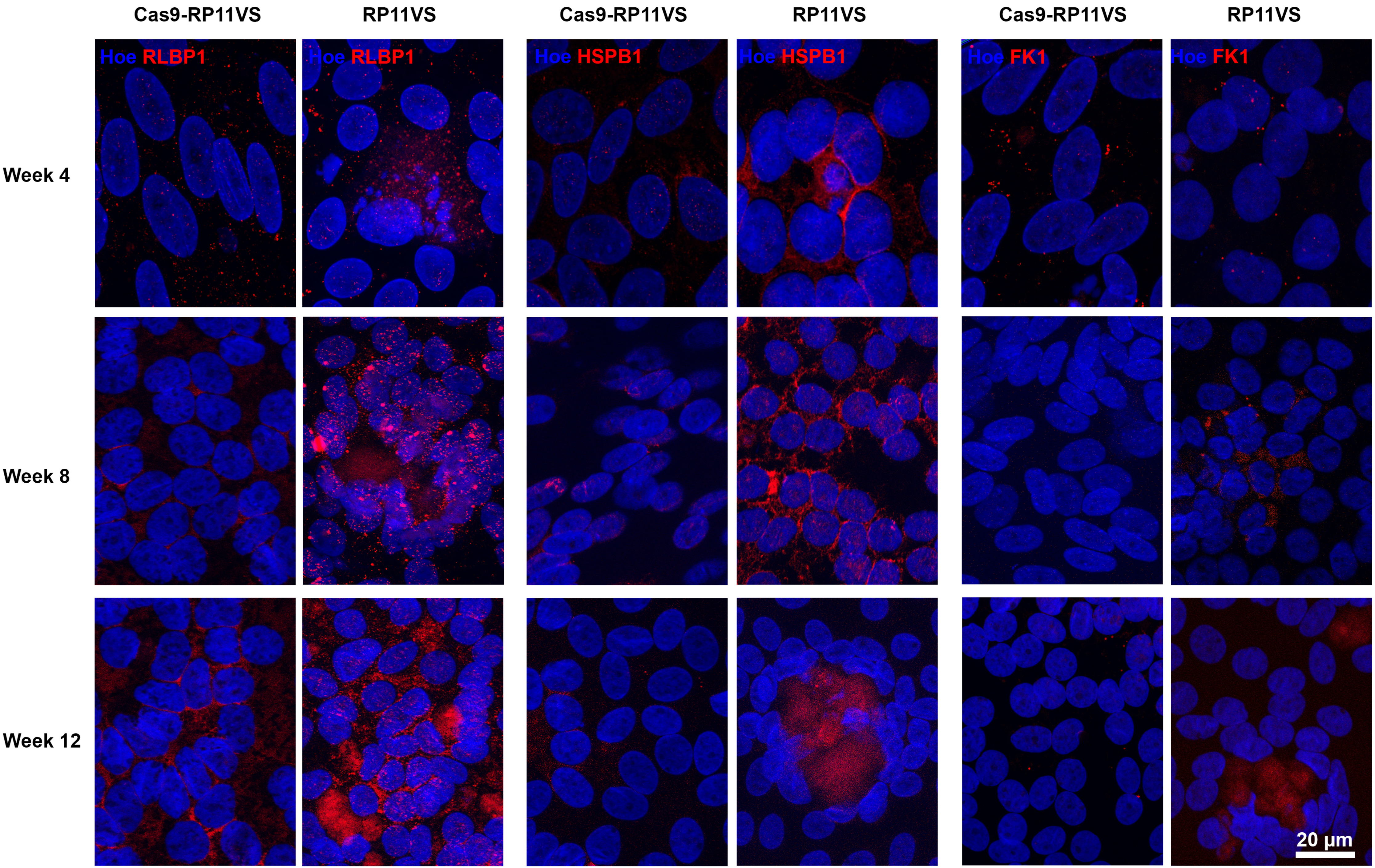

### Supplementary Figure 6

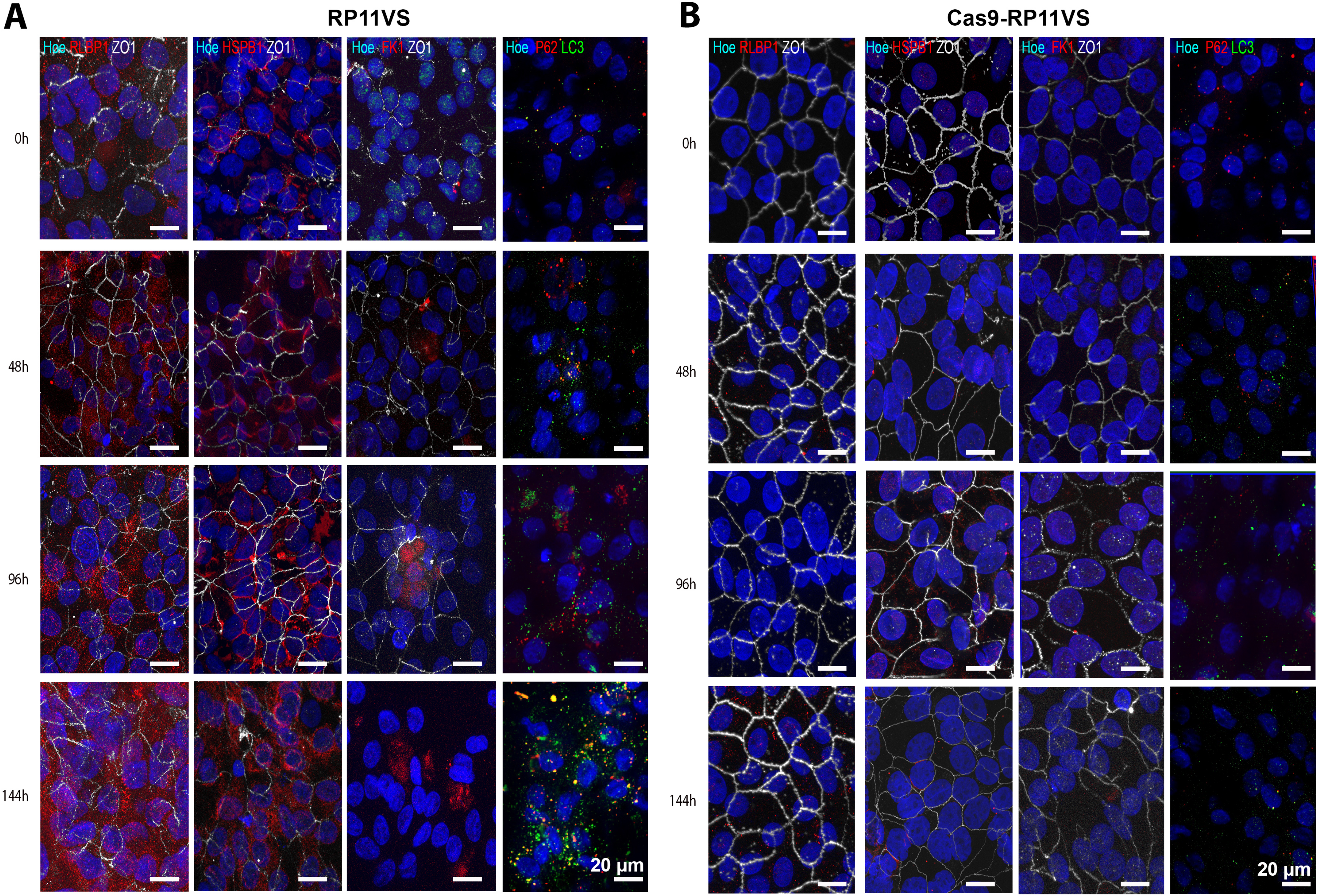

### Supplementary Figure 7

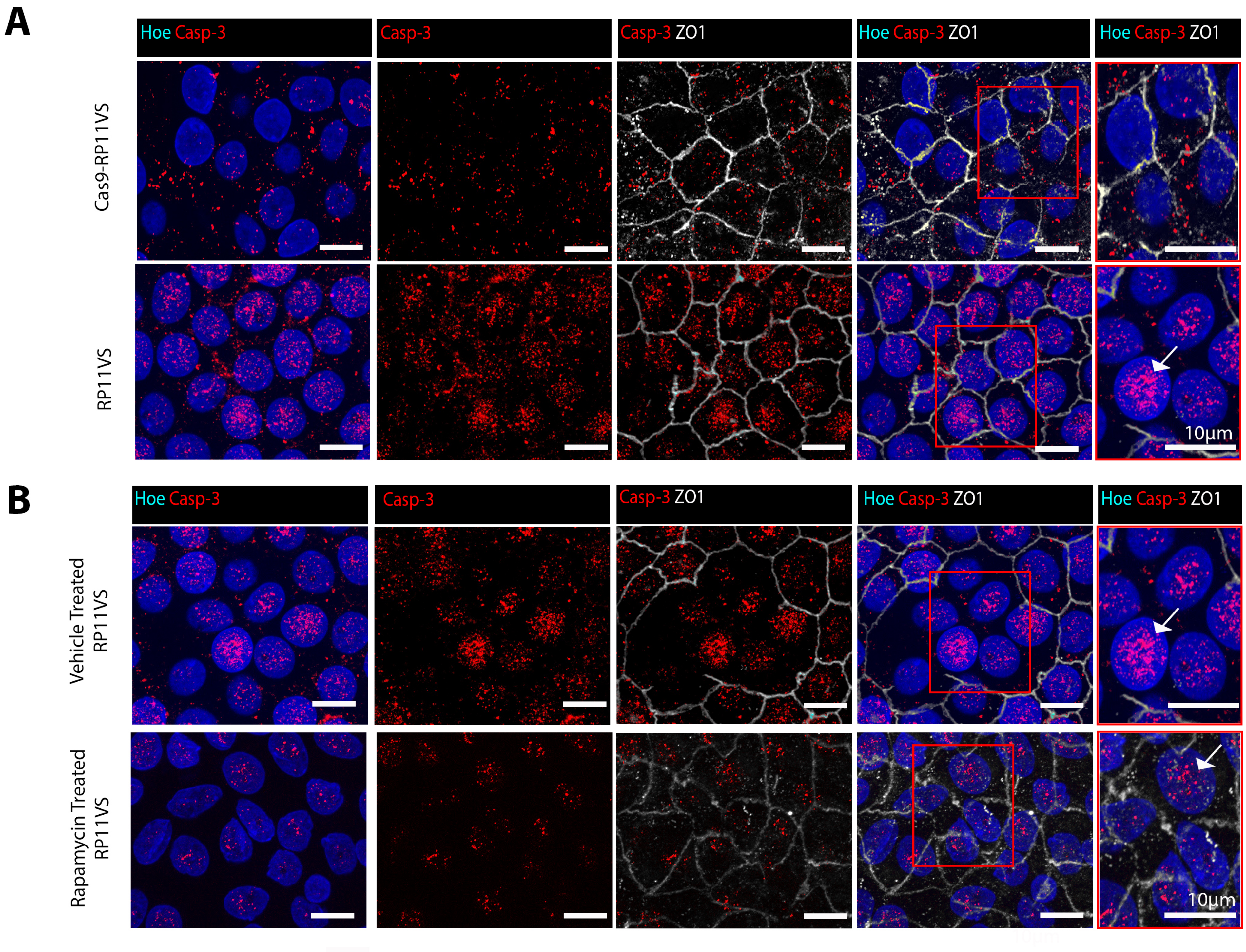

### Supplementary Figure 8

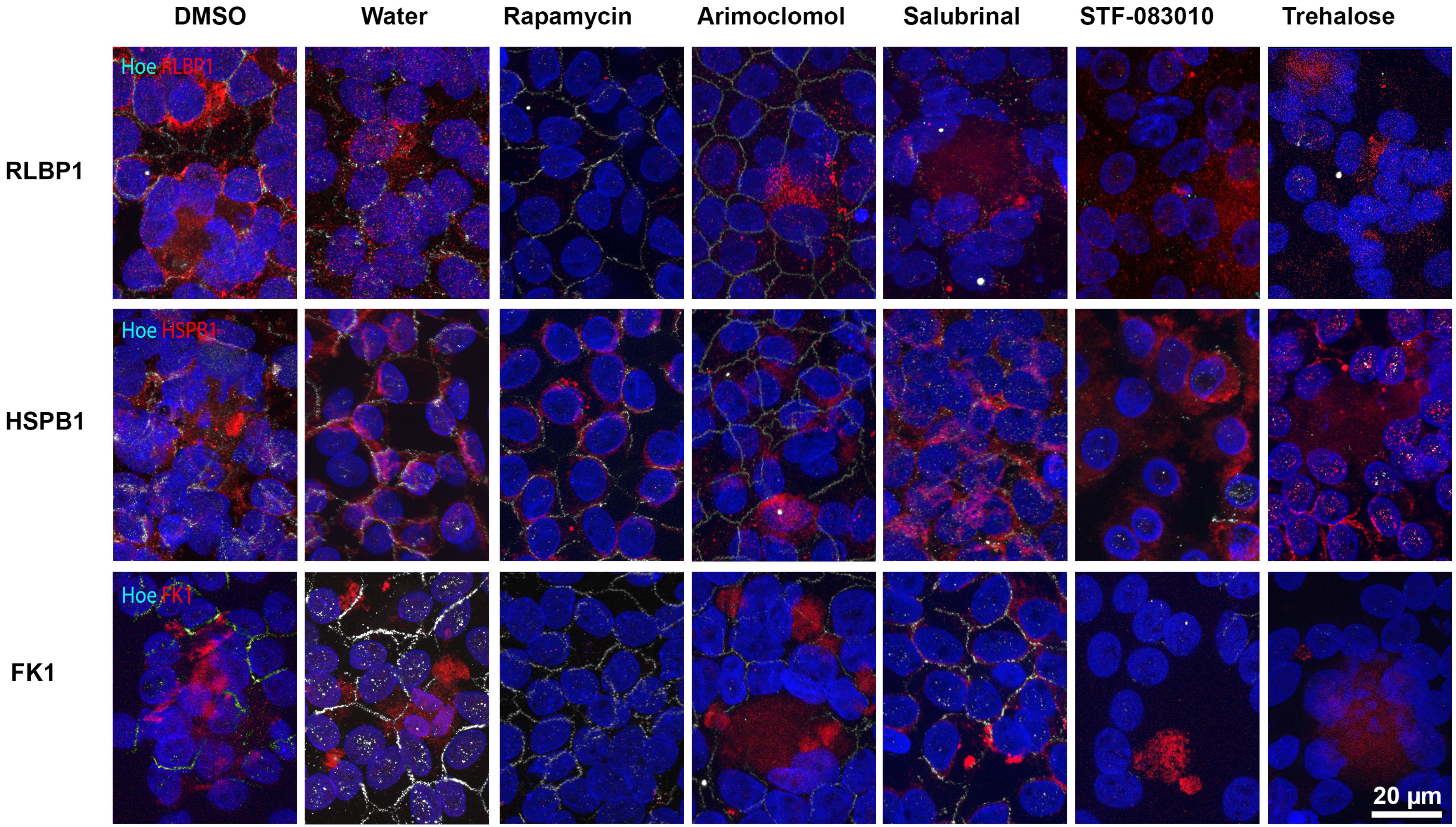
