## Supplementary Table 1 for "Progressive protein aggregation in PRPF31 patient retinal pigment epithelium cells: the mechanism and its reversal through activation of autophagy"

| **Antibody** | **Host** | **Source** | **Cat. No.** | **Dilution** |
| --- | --- | --- | --- | --- |
| ACTB | Mouse | Santa Cruz Biotechnology | sc-47778 | WB: 1:500 |
| ATG5 | Rabbit | Sigma-Aldrich | A0856 | WB: 1:500 |
| Beclin 1 | Rabbit | Abcame | ab210498 | WB: 1:500 |
| Caspase-3 | Rabbit | Cell Signaling Technology | 9661S | IF: 1:400 |
| Coilin | Mouse | Santa Cruz Biotechnology | sc-55594 | IF: 1:200 |
| FK1 | Mouse | Enzo Life Sciences | PW8805 | IF: 1:500  WB: 1:500 |
| Fus | Rabbit | Proteintech | 11570-1-AP | IF: 1:50  WB: 1:500 |
| GAPDH | Mouse | Santa Cruz Biotechnology | sc-47724 | WB: 1:500 |
| HSPA2 | Mouse | Antibodies-online GmbH | ABIN561382 | IF: 1:100  WB: 1:500 |
| HSPA4L | Rabbit | Novus Biologicals | NBP2-48700 | IF: 1:250  WB: 1:500 |
| HSPB1 | Mouse | Insight Biotechnology | OASG03654 | IF: 1:250  WB: 1:500 |
| LC3B | Rabbit | Cell Signaling Technology | 3868S | IF: 1:500  WB: 1:500 |
| MAP1LC3B | Rabbit | Cell Signaling Technology | 3868S | IF: 1:200  WB: 1:500 |
| p62 | Mouse | BD Biosciences | 610832 | IF: 1:200  WB: 1:500 |
| Phosphor-eLF2a | Rabbit | Cell Signaling Technology | 3597S | WB: 1:1000 |
| PRPF31 | Goat | Sigma | SAB2500828 | IF: 1:250  WB: 1:500 |
| p-S6 (Ser235/236) | Rabbit | Cell Signaling Technology | 4858S | WB: 1:500 |
| 20S proteasome α/β subunits | Rabbit | Enzo Life Sciences | BML-PW8155-0025 | WB: 1:1000 |
| RLBP1 | Mouse | Abcam | ab15051 | IF: 1:100  WB: 1:500 |
| SC-35 | Mouse | Santa Cruz Biotechnology | sc-53518 | IF: 1:100 |
| SF3B1 | Mouse | Santa Cruz Biotechnology | sc-514655 | IF: 1:150 |
| S6 | Rabbit | Cell Signaling Technology | 2217S | WB: 1:1000 |
| ZO1 | Goat | St John's Laboratory | STJ140055 | IF: 1:50 |
| ZO1 | Rabbit | Invitrogen | 61-7300 | IF: 1:50 |
| ZO1 | Mouse | Thermo Scientific | 33-9100 | IF: 1:50 |

**Table S1: Summary of antibodies used in this study.**

**Primary antibodies**

| **Antibody** | **Source** | **Cat. No.** | **Dilution** |
| --- | --- | --- | --- |
| Donkey anti-Goat Alexa 488 | Life Technologies | A11055 | 1:800 |
| Donkey anti-Goat Alexa 647 | ThermoFisher | A21447 | 1:800 |
| Donkey anti -Mouse FITC | Jackson ImmunoResearch | 715-095-151-JIR | 1:800 |
| Donkey anti-Mouse Alexa 488 | Life Technologies | A21202 | 1:800 |
| Donkey anti-Rabbit Alexa 546 | Life Technologies | A10040 | 1:800 |
| Polyclonal Swine Anti-Rabbit Immunoglobulins/HRP, | Agilent Dako | P0399 | 1:2000 |
| Rabbit Anti-Mouse Immunoglobulins/HRP | Agilent Dako | P0260 | 1:2000 |
| Polyclonal Rabbit Anti-Goat Immunoglobulins/HRP, | Agilent Dako | P0449 | 1:2000 |

**Secondary antibodies**
